# Serial Lift-Out – Sampling the Molecular Anatomy of Whole Organisms

**DOI:** 10.1101/2023.04.28.538734

**Authors:** Oda Helene Schiøtz, Christoph J.O. Kaiser, Sven Klumpe, Dustin R. Morado, Matthias Poege, Jonathan Schneider, Florian Beck, Christopher Thompson, M. Jürgen Plitzko

**Affiliations:** Research Group CryoEM Technology, Max Planck Institute of Biochemistry, Martinsried, Germany; Department Cell and Virus Structure, Max Planck Institute of Biochemistry, Martinsried, Germany; Department Molecular Structural Biology, Max Planck Institute of Biochemistry, Martinsried, Germany; Materials & Structural Analysis, Thermo Fisher Scientific, Eindhoven, The Netherlands

**Keywords:** Cryo-focused ion beam (cryo-FIB), Lift-Out, C. elegans, cryo-electron tomography (cryo-ET)

## Abstract

Cryo-focused ion beam milling of frozen-hydrated cells and subsequent cryo-electron tomography (cryo-ET) has enabled the structural elucidation of macromolecular complexes directly inside cells. Application of the technique to multicellular organisms and tissues, however, is still limited by sample preparation. While high-pressure freezing enables the vitrification of thicker samples, it prolongs subsequent preparation due to increased thinning times and the need for extraction procedures. Additionally, thinning removes large portions of the specimen, restricting the imageable volume to the thickness of the final lamella, typically < 300 nm. Here, we introduce Serial Lift-Out, an enhanced lift-out technique that increases throughput and obtainable contextual information by preparing multiple sections from single transfers. We apply Serial Lift-Out to *C. elegans* L1 larvae yielding a cryo-ET dataset sampling the worm’s anterior-posterior axis and resolve its ribosome structure to 7 Å, illustrating how Serial Lift-Out enables the study of multicellular molecular anatomy.

## Introduction

Single particle analysis (SPA) by cryo-transmission electron microscopy (cryo-TEM) has become a key technique to study the structure of isolated biological macromolecules at high-resolution (Kühlbrandt 2014). While SPA routinely reaches resolutions where protein side-chains can be fitted unambiguously, the reductionist approach of studying protein complexes *in vitro* loses all information concerning their molecular sociology: the interaction of the molecular complexes in their natural environment (Beck & Baumeister 2016). Conversely, *in situ* cryo-electron tomography (cryo-ET) allows for the reconstruction of pleomorphic structures such as the crowded environment of the cell at molecular resolution, maintaining the interaction and localization of protein complexes within the biological system (Dietrich et al 2022, Gupta et al 2021, Hoffmann et al 2022, O’Reilly et al 2020, Plitzko et al 2017, Watanabe et al 2020).

One of the primary factors limiting the resolution of cryo-TEM is inelastic scattering. As the mean free path of an electron in vitrified biological samples is about 300-400 nm, samples beyond the size of viruses and small prokaryotic cells are generally too thick for cryo-ET (Liedtke et al 2022, O’Reilly et al 2020). In consequence, two main sample thinning methods to obtain electron-transparent specimens for cryo-ET have been developed: cryo-ultramicrotomy and cryo-FIB milling. Cryo-ultramicrotomy encompasses the thin-sectioning of vitreous cells and tissues with a diamond knife (Al-Amoudi et al 2004). The shearing forces at the knife’s edge, however, cause mechanical artifacts such as crevices and compression in the resulting sections (Al-Amoudi et al 2003). More recently, the focused ion beam (FIB) instrument has been widely adopted for sample thinning at cryogenic temperatures. While not completely damage-free (Berger et al 2023, Lucas & Grigorieff 2023), the technique bypasses the mechanical artifacts of cryo-ultramicrotomy (Marko et al 2007, Rigort et al 2012) and has been shown to yield data that can allow for the elucidation of ribosomes to side-chain resolution (Hoffmann et al 2022). The automation of lamella preparation by cryo-FIB milling has also reduced the need for user expertise and manual intervention (Buckley et al 2020, Klumpe et al 2021, Zachs et al 2020).

Prior to thinning, the sample must be cryogenically fixed by cooling at a sufficiently high rate to prevent ice crystal formation, resulting in a vitrified sample. There are two main methods available for vitrification: plunge freezing, in which the sample is immersed at ambient pressure into liquid ethane or ethane-propane mixture (Tivol et al 2008), and high pressure freezing (HPF), in which the sample is cooled with a jet of liquid nitrogen at a pressure of ∼2000 bar. While the former yields samples that are easily FIB-milled, the sample thickness that can reliably be vitrified is limited to roughly 10 µm. HPF, on the other hand, allows for the vitrification of samples up to a thickness of roughly 200 µm (Dubochet 1995).

Consequently, HPF greatly expands the size range of biological samples that can be vitrified but comes at the cost of embedding the specimen in a thick layer of ice defined by the depth of the freezing receptacle. This increased sample thickness leads to longer milling times. While samples up to a thickness of about 50 µm can be prepared by milling lamellae directly on the grid following the ‘waffle’ method (Kelley et al 2022), lamellae from thicker samples have, to date, only been prepared by cryo-lift-out (Klumpe et al 2021, Kuba et al 2021, Mahamid et al 2015, Parmenter et al 2016, Rubino et al 2012, Schaffer et al 2019).

Cryo-lift-out refers to the extraction of the material for lamella preparation from bulk HPF sample and subsequent transfer and attachment to a lift-out receiver grid, conventionally a half-moon shaped grid (Giannuzzi et al 2005). Two main types of micromanipulator devices are currently available for cryo-lift-out: sharp needles and a cryo-gripper (Klumpe et al 2022). Initially, trenches are milled around the area of interest leaving it connected to the bulk material on a single side. For specimens in HPF sample carriers, the material must additionally be cleared from below. After these preparatory steps, the lift-out device is brought in contact with the volume to be extracted. The remaining connection to the bulk material is removed, the micromanipulator is used to transfer the extracted volume and redeposition milling (Schreiber et al 2018) is used to attach the volume to the receiver grid. Finally, an electron transparent lamella is prepared (Parmenter & Nizamudeen 2021).

While widely used in materials science at room temperature (Giannuzzi et al 2005), cryo-lift-out of biological samples has remained primarily proof-of-concept (Klumpe et al 2021, Parmenter et al 2016, Rubino et al 2012, Schaffer et al 2019, Schreiber et al 2018). This is due to a number of factors, e.g. the need for cooled micromanipulator devices and accompanying workflow adaptations required resulting in limited throughput and problems with lamella loss during transfer to the TEM (Schaffer et al 2019).

Another limitation to cryo-lift-out, as well as on-grid lamella preparation is the loss of contextual information. Only a fraction of the sample volume (∼1% for larger eukaryotic cells, < 1% for multicellular specimens) ends up inside the final lamella for cryo-ET data acquisition. Techniques exist that are capable of capturing larger volumetric data and tracking morphology at comparatively large volume scales. Examples of such techniques are X-ray tomography and various volume electron microscopy (vEM) techniques: serial FIB milling and scanning electron microscopy, and serial sectioning of plastic embedded samples imaged by TEM or STEM. These techniques, however, currently cannot achieve the resolution attainable by cryo-ET at an equivalent sample preservation state due to physical limitations in imaging or the necessity of fixation and contrasting (Dumoux et al 2023, Spehner et al 2020).

In this work, we describe a novel cryo-lift-out approach that creates a series of lamellae from one lift-out volume that we term Serial Lift-Out. Inspired by diamond knife serial sectioning, Serial Lift-Out retains more contextual information than previous procedures and increases the throughput of cryo-lift-out by an order of magnitude. It sets the stage for the study of multicellular organisms and tissues by cryo-ET – applications that previously seemed practically impossible (Parmenter et al 2016). We demonstrate Serial Lift-Out on high-pressure frozen *Caenorhabditis elegans* L1 stage larvae, sampling their ultrastructure along the anterior-posterior axis by cryo-electron tomography. From the resulting dataset, we reconstruct the nematode’s ribosome to 7 Å resolution by subtomogram averaging, exemplifying the enormous potential of Serial Lift-Out for the study of the molecular anatomy of multicellular systems.

## Results

### The concept behind Serial Lift-Out

The most time-consuming steps in cryo-lift-out are the preparation of the extraction sites and the transfer of the extracted volume to the receiving grid. Each repetition of the lift-out cycle therefore adds significant time investment with low yield. Scaling up the extracted volume and producing multiple lamellae from a single lift-out bypasses this repetitive, time-consuming trench milling and transfer, increases throughput and, foremost, provides more contextual information about the targeted material.

A specific implementation of this concept is the extraction of an entire L1 larva of *C. elegans*, followed by repeated steps of attachment to the receiving grid, sectioning and transfer of the remaining volume to the next attachment site (Figure 1). While material is still lost during sectioning and thinning, a large fraction of the worm is made readily accessible to cryo-TEM data acquisition.

**Figure 1:**
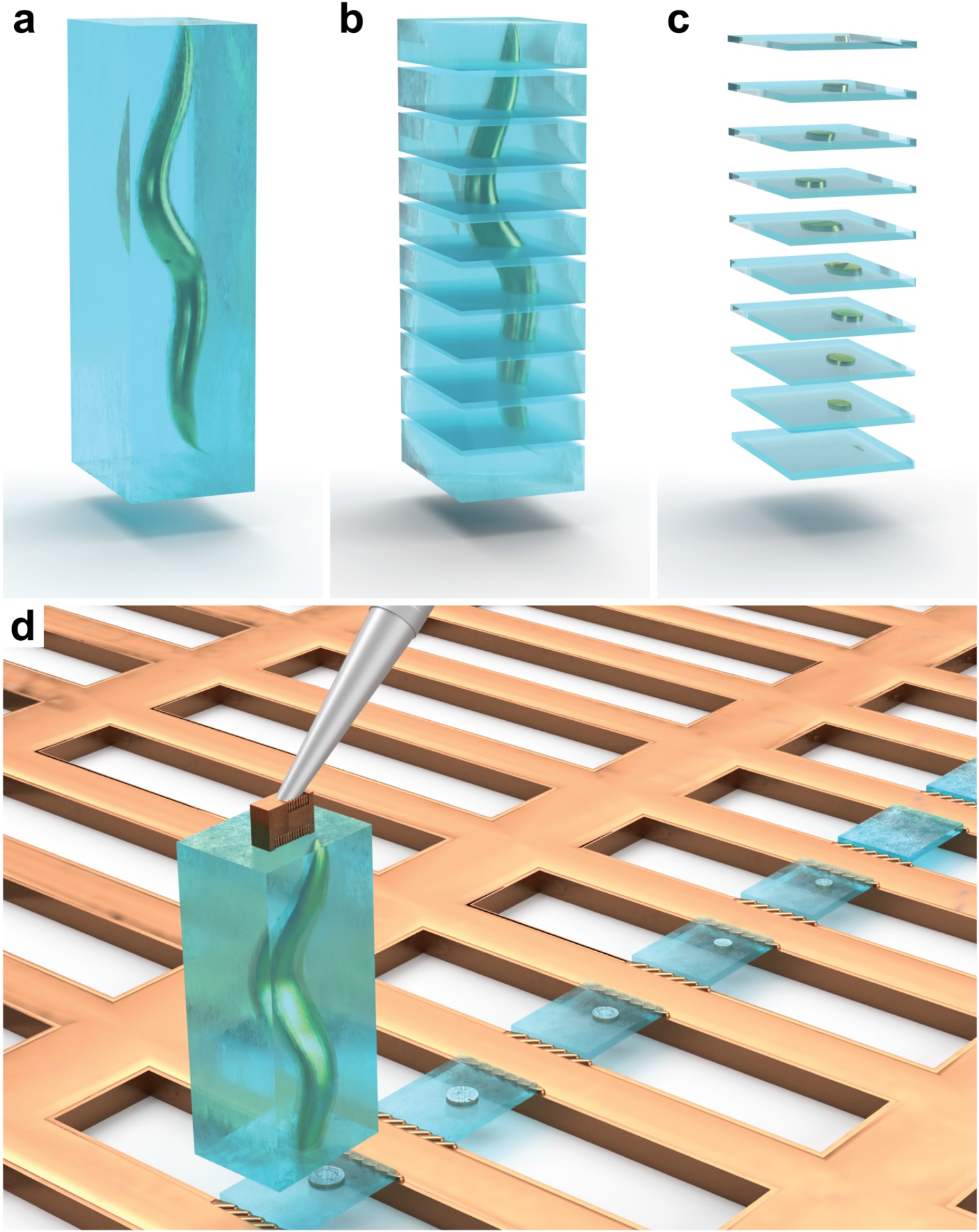
A schematic of Serial Lift-Out. An illustration of the Serial Lift-Out method, exemplified by the process being performed on a *C. elegans* L1 larva embedded in an extracted volume of vitrified ice (blue). **a**, The extracted volume containing the larva. **b, c**, Schematic representation of the resulting Serial Lift-Out sections and lamellae, respectively. **d**, Illustration of the double-sided attachment Serial Lift-Out procedure. The extracted volume shown in panel **a** is attached to the lift-out needle via a copper block adaptor and transferred to a rectangular mesh receiver grid. Several sections are shown, obtained by the repetition of the attachment of the bottom part of the volume to the grid bars and subsequent sectioning.

Previous lift-out approaches extracted volumes approximately the size of the final lamella (Figure 1 - Figure Supplement 1a), with some excess material for stability during transfer and attachment. In order to section multiple lamellae from a single cryo-lift-out transfer, the volume extracted by cryo-lift-out needed to be increased. To ease the manipulation of such large volumes, we introduced a copper block that acts as an adapter between the needle and the extracted volume. This copper block is created from the receiver grid and attached to the needle before lift-out (Figure 1 - Figure Supplement 2, Figure 1 – Figure Supplement 3). As copper has higher redeposition rates than the tungsten needle, the copper block adapter results in significantly more resilient “welding” of the specimen to the micromanipulator.

Another volume limiting factor is the ablation rate of the ion beam. Most cryo-FIB machines are equipped with a gallium ion source rendering it impractical to mill beyond 50 µm in depth. In addition, for samples frozen in HPF carriers, preparation requires an undercut, the removal of material below the extracted volume, to detach the extraction volume from the bulk. In combination, these factors generally result in maximum extraction volumes of approximately 20 µm × 20 µm × 10-30 µm (length × width × height, Figure 1 - Figure Supplement 1b).

In order to extract larger volumes, we performed lift-out on an HPF ‘waffle’-type sample. HPF ‘waffle’ samples are prepared by freezing sample on a grid that is sandwiched between HPF carriers (Kelley et al 2022). The final thickness is therefore defined by the type of grid and spacer being used during freezing. For a 25 µm thick specimen, extraction volumes of up to 200 µm × 40 µm × 25 µm can easily be obtained by performing lift-out with the sample surface oriented perpendicular to the ion beam (Figure 1 – Figure Supplement 1c). The same orientation is used during trench milling and will be referred to as ‘trench milling orientation’ (Figure 1 – Figure Supplement 3c,f). Such large volumes extracted at the trench milling orientation can yield many sections from a single lift-out which can be subsequently thinned to lamellae for cryo-ET data acquisition (Figure 1b,c).

### Application of Serial Lift-Out to *C. elegans* L1 larvae

To assess the feasibility of obtaining serial lamellae from a single lift-out transfer, we performed the procedure on *C. elegans* L1 larvae that had been vitrified using a modified protocol of the ‘waffle’ method (Figure 2). The sample contained many L1 larvae embedded in an approximately 25 µm thick layer of ice (Figure 2 - Figure Supplement 1). Two sites from two different grids were selected for preparation by correlating cryo-fluorescence light microscopy with SEM and FIB view images. Following the geometry described in Figure 1 - Supplement 1c, the sites were prepared by milling trenches around the larva, yielding a 30 µm × 110 µm × 25 µm and a 40 µm × 180 µm × 25 µm extraction volume (Figure 2a, Figure 2 – Figure Supplement 3a). Two different attachment strategies were explored, with resulting lamellae either being attached on one side (single-sided attachment) or two sides (double-sided attachment). To extract the volumes, the needle with the copper adapter (Figure 2a, Figure 2 – Figure Supplement 3b) was attached to the sample using redeposition: copper material redepositing onto the surface of the extraction volume during milling. The extraction volume was subsequently milled free from the bulk material and lifted out.

**Figure 2:**
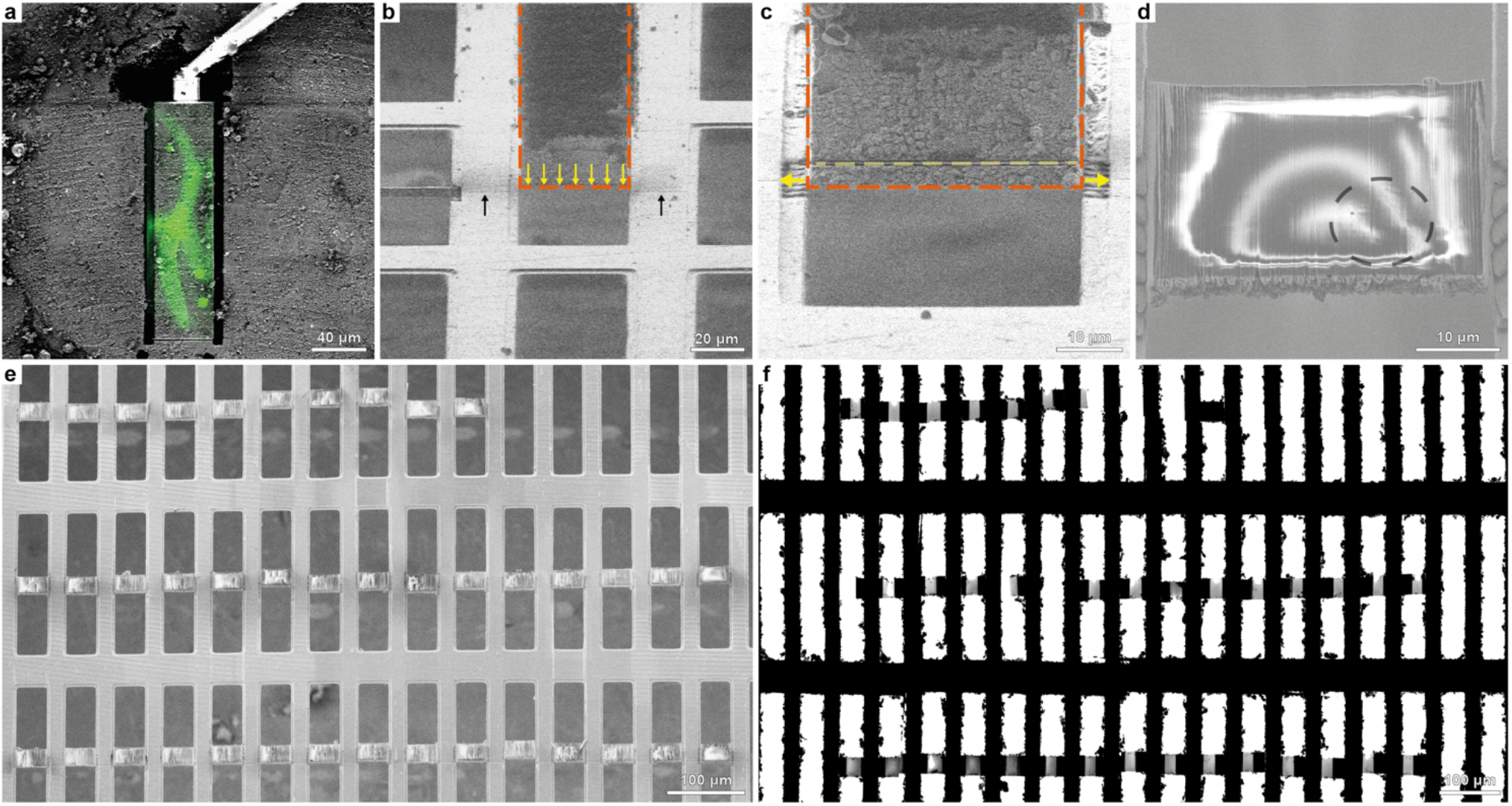
A workflow for double-sided attachment Serial Lift-Out. **a**, FIB image of the prepared extraction site with overlaid correlated fluorescence data (green) indicating the larva being targeted. The micromanipulator is attached to the extraction volume by redeposition from the copper adapter (trench milling orientation). **b**, The extracted volume (orange dashed line) is lowered into position between two grid bars in lamella milling orientation. The lower front edge of the volume (yellow arrows) is aligned to the pre-milled line mark (black arrows). **c**, Double-sided attachment by redeposition from the grid bars (yellow arrows indicate direction of milling), followed by line pattern milling releasing the section of a desired thickness (dashed yellow line). Orange dashed line indicates the outline of the extracted volume. **d**, SEM image of a typical section after being released from the extracted volume. Black dashed line indicates the outline of the worm cross-section. **e**,**f**, SEM (**e**) and TEM (**f**) overview images of the 40 double-sided attached sections obtained. Figure 2 - Movie Supplement 1 summarizes the process.

The volume was then transferred to the receiver grid. For single-sided attachment, a modified grid based on customizing a standard 100 mesh copper grid (Figure 2 - Figure Supplement 2) was used. Removing every second row of grid bars yielded an array of trimmed grid bars that resemble the pins of a standard half-moon shaped lift-out grid (Figure 2 - Figure Supplement 2e). Alternatively, a copper grid with rectangular meshes can be used for double-sided attachment of volumes roughly as wide as the mesh (∼40 µm for the 400/100 rectangular mesh grids used here, Figure 2b). Double-sided attachment has the added benefit of increased section stability by eliminating the free-standing side (Figure 2 – Figure Supplement 3f).

The volume’s lower front edge was precisely aligned to the front edge of the attachment pin (Figure 2 – Figure Supplement 3c) or a previously prepared alignment line pattern milled onto the receiver grid (Figure 2b). Then, redeposition milling was used to attach the lower part of the volume to the grid bar(s) (Figure 2c, Figure 2 – Figure Supplement 3d, yellow arrows). After attachment, the lower part of the target material was separated from the extracted volume using a line pattern (Figure 2c, Figure 2 – Figure Supplement 3d, yellow dashed line), leaving a ∼4 µm thick section (Figure 2d, Figure 2 – Figure Supplement 3e,f). Once sectioned, the remaining volume attached to the lift-out needle was transferred to the next attachment site. This procedure was iterated until no material remained. As a result, many sections were produced from a single cryo-lift-out transfer (Figure 2e-f). To assess the general applicability of the approach to non-’waffle’ type samples, an additional experiment was performed on *D. melanogaster* egg chambers, high-pressure frozen in standard sample carriers. From a ∼20 µm long and ∼15 µm deep extracted volume we produced sections of ∼1-2 µm thickness (Figure 2 - Figure Supplement 4).

### Sampling organismal cellular anatomy along the *C. elegans* L1 larva at molecular resolution

We prepared dozens of sections along the anterior-posterior axis of *C. elegans* L1 larvae: 12 single-sided attached and 40 double-sided attached lamellae (Figure 2 – Figure Supplement 3g, Figure 2e). After transfer to the TEM, 8 out of 12 and 32 out of 40 lamellae were recovered, respectively (Figure 3 – Figure Supplement 1a, Figure 3 – Figure Supplement 2a). Lamella loss during transfer is common due to manual grid handling steps. The increased rate of successfully transferred lamellae from 66% for single-sided to 80% for double-sided attachment is indicative of the increased lamella stability of double-sided attachment.

**Figure 3:**
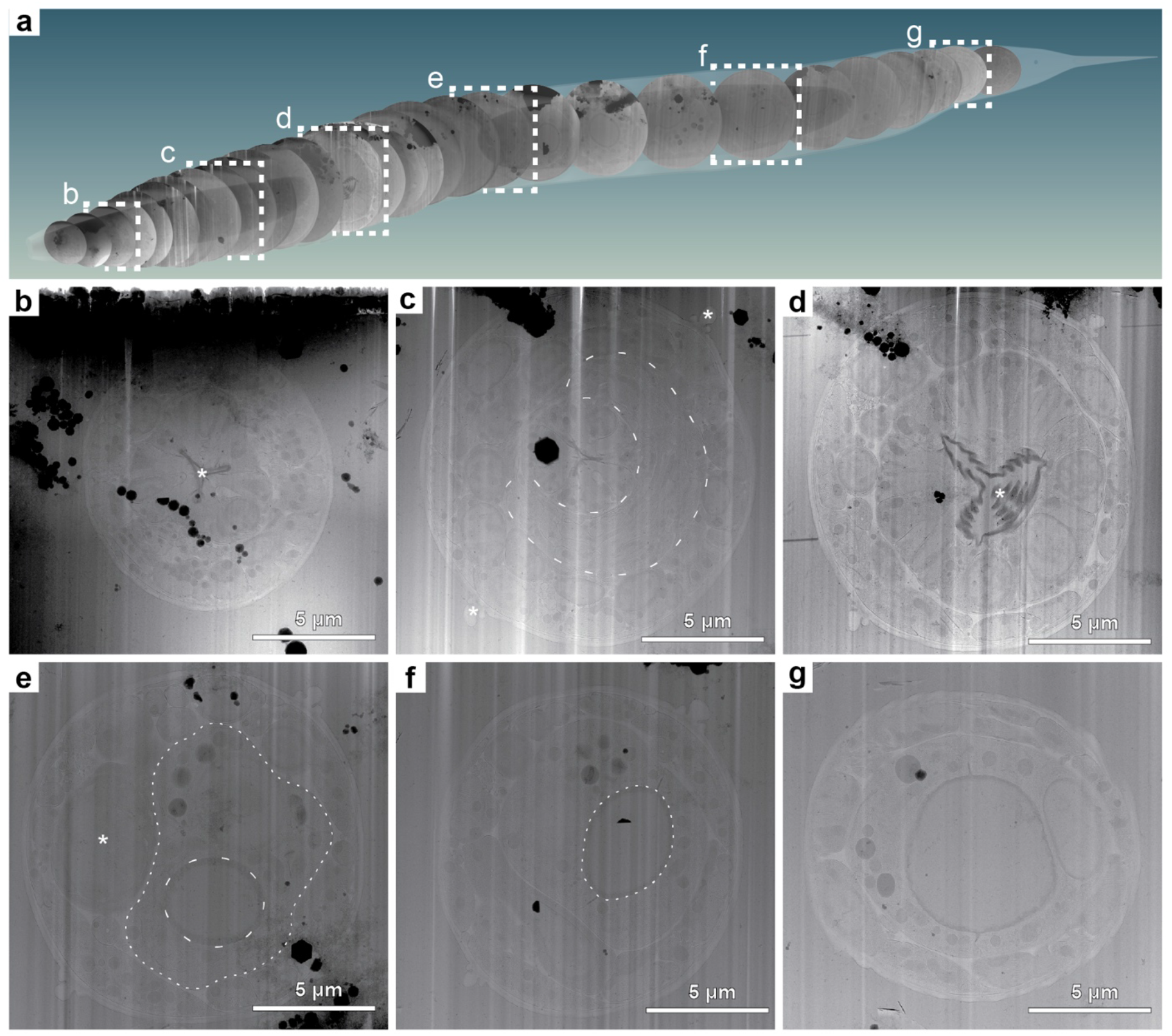
Lamella TEM overviews sample the anatomy of a *C. elegans* L1 larva along the anterior-posterior body axis. Native tissue scattering contrast is sufficient to extract a considerable amount of anatomical information from low magnification TEM overviews of the lamellae generated in the double-sided attachment Serial Lift-Out experiment. **a**, Schematic representation of 29 body transverse section overviews obtained from the final lamellae along the anterior-posterior body axis of an L1 larva. Anterior is located to the left, posterior to the right. The OpenWorm project model of the adult *C. elegans* worm was used for illustrating the possibility of back mapping since a cellular model of the L1 larva only exists for its head and not the entire body (Britz et al 2021). The cross-sections were cropped from lamella overview images and mapped back to a position derived from the known sectioning distance and anatomical features discernible in the corresponding cross-section. The dashed white frames indicate overviews with corresponding magnified representations in panels **b**-**g. b**, This lamella originates from ∼15 µm along the anterior-posterior axis. Clearly visible are the three lobes of the anterior pharyngeal lumen in the center of the worm cross-section (asterisk) and the relatively electron dense pharyngeal lining. **c**, Overview from the anterior part of the pharyngeal isthmus ∼42 µm along the anterior-posterior axis. Note the nerve ring (dashed line) surrounding the central pharynx. Additionally, the alae (asterisks) running along the left and right lateral side of the worm become obvious. **d**, Overview of a lamella of roughly the center of the posterior pharyngeal bulb region. The central grinder organ is clearly discernible (asterisk). This section is positioned ∼65 µm along the anterior-posterior axis. **e**, A section roughly mid-body. The intestinal lumen (dashed line) and intestinal cells (dotted line) are obvious. The darker cell slightly left of the body center is likely one of the gonadal primordial cells (asterisk). The section is from ∼115 µm along the anterior-posterior axis. **f**, In this mid-body section, the intestinal lumen (dashed line) can again be clearly discerned. The section can be mapped to ∼132 µm along the anterior-posterior axis. **g**, Section showing the intestinal lumen at ∼155 µm along the anterior-posterior axis.

From these successfully transferred lamellae, overview maps were recorded (approximately 20 µm × 25 µm in size, Figure 3 – Figure Supplement 1, Figure 3 – Figure Supplement 2) depicting clear larval cross-sections. These overviews allowed the identification and assignment of anatomical structures and tissues such as the pharynx, the body-wall muscle cells, neurons, the hypodermis and seam cells. Larger organelles such as nuclei, mitochondria, Golgi cisternae, storage granules, junctional regions, bundles of actin filaments and microtubules were clearly discernible (Figure 3b-g, Figure3 - Movie Supplement 1). Given the stereotypic body plan of *C. elegans*, the information obtained from the overviews together with the sectioning thickness was used to determine the approximate location of the sections in the worm, as schematically shown in Figure 3a.

The biological area within each lamella can, due to the circular nature of cross sections, be estimated by ?r^2^, where the average radius is around 5 µm, resulting in a mean of 78.5 µm^2^ imageable area per lamella. For the single-sided attachment experiment, the total imageable area was ∼ 630 µm^2^ and allowed for the collection of 57 tilt series, each with a field of view of ∼1.2 µm in size. The double-sided attachment experiment yielded ∼2500 µm^2^ imageable area from which a total of 1012 tilt series with a field of view of ∼750 nm in size were collected. The double-sided attachment experiment allowed us to sample a large fraction of all tissues and cell types along the anterior-posterior body axis of the *C. elegans* L1 larva.

To illustrate the information level already present at intermediate magnification (11,500x), we manually segmented a representative cross-section (Figure 4A). The segmentation illustrates that the cuticle clearly delineates the body cross-section. Body-wall muscle cells are obvious due to their pronounced distal actomyosin pattern. The pharyngeal lumen’s trilobal structure demarcates the body center and is surrounded by the three alternating pharyngeal muscle and marginal cells. Pharyngeal neurons and gland cell process cross-sections are embedded in the pharyngeal muscle cells. Nuclei are easy to discern due to their ribosome-decorated double-membrane and their denser, less granular interior. Neurons in general can be discerned by their appearance as round or tubular cells, often grouped in bundles. These clearly interpretable overviews allow targeting of tissue-specific cellular structures or cellular protein complexes with known location (e.g. sarcomeric proteins).

**Figure 4:**
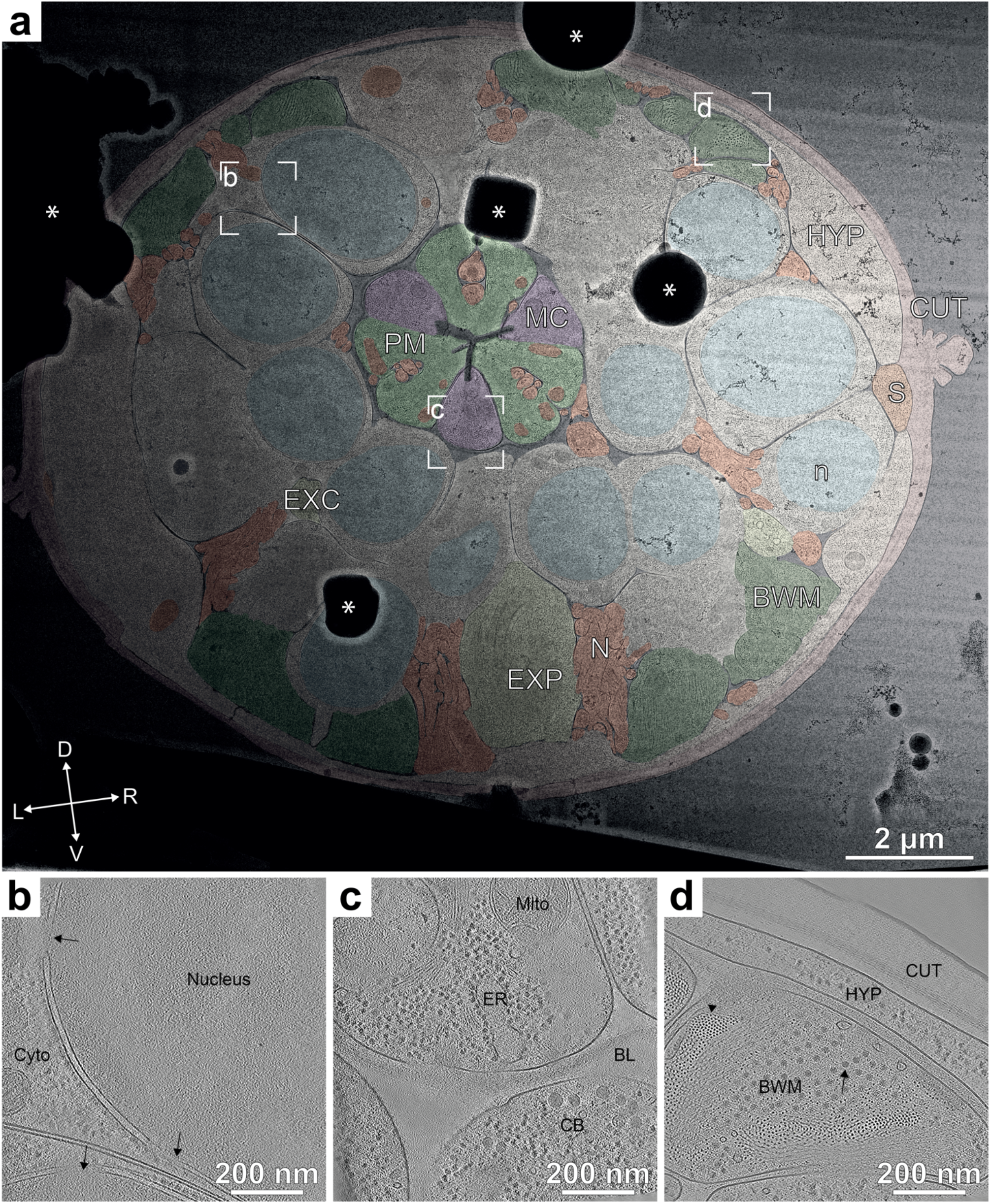
Representative overview and tomograms of the L1 larval pharyngeal isthmus region. **a**, Overview montage (magnification 11,500x) of the head region lamella. This section is located within the pharyngeal isthmus just posterior to the nerve ring. Cell types are colored according to the WormAtlas color code (Altun et al 2002-2023), for nuclei and mitochondria, arbitrary colors were chosen (HYP: hypodermis, S: seam cells, PM: pharyngeal muscle, MC: marginal cells, EXP: excretory pore, EXC: excretory canal, N: neuronal tissue, CUT: cuticle, n: nucleus. The white dashed rectangles show the positions of the tilt series acquired and corresponding reconstructions shown in **b**-**d**. Asterisks indicate ice contamination. The cross indicates the dorsal(D)-ventral(V) and left(L)-right(R) body axes. **b**, Tomographic slice of the perinuclear region of most likely a neuronal cell. The nucleus exhibits a different granularity and density than the cytoplasm (Cyto), from which it is separated by the nuclear envelope, which in turn is heavily decorated with ribosomes and contains nuclear pores (arrows). **c**, Tomographic slice of the pharynx. Central in the upper half is a marginal cell containing endoplasmatic reticulum (ER) and mitochondria (Mito). A neighboring neuronal cell body (CB) is separated from the pharynx by a diffuse density, which is the pharyngeal basal lamina (BL). **d**, Tomographic slice of a body wall muscle cross section (BWM). Clearly visible are the actin filaments in top view (arrowhead) surrounding a bundle of thick filaments (arrow). The thick filaments are partially interspersed with actin filaments. The muscle cell neighbors a hypodermal cell (HYP), from which it is separated by a space filled with a diffuse density, likely the body-wall muscle basal lamina. The next and last layer of the larval body wall is the cuticle (CUT). Most notably within this chitinous/collagenous structure, a fence-like array of denser structures can be discerned, likely cuticular collagen.

### Quality assessment and subtomogram analysis

To assess the quality of lamella thinning, tomogram thickness was measured for all 57 tomograms from the single-sided attachment and a random subset of 132 tomograms for the double-sided attachment experiment. The thickness for single-sided attachment was 253 nm ± 125 nm. The thickness of the tomograms from the double-sided attachment experiment was more uniform, 303 ± 40 nm (Figure 4 – Figure Supplement 1). The broader thickness distribution in single-sided attachment most likely stems from lamella bending and movement during milling due to the free-standing edge of the lamella.

The tilt series we obtained allowed us to reconstruct the 3D cellular architecture of cell types in different tissues such as the neuronal nuclear periphery (Figure 4b), pharyngeal marginal cells and neuronal cell bodies (Figure 4c) and body wall muscle, the hypodermis and the collagenous network of the cuticle (Figure 4d).

The sectioning plane obtained when doing Serial Lift-Out from ‘waffle’-type samples provides worm cross-sections. For a number of structures, such as nuclei, the difference between oblique longitudinal and transverse sectioning is minimal (Figure 4a, Figure 4 – Figure Supplement 2). When investigating anisotropic structures such as the body wall muscle or pharynx, however, sectioning direction can greatly impact the interpretation of higher order structure. In longitudinal sectioned sarcomeres, located in body-wall muscle, appear as filaments running along the image plane. These filaments show an orientational distribution strongly biased towards the side view. In contrast, tomograms from Serial Lift-Out show transverse sections of the body-wall muscle. In this sectioning plane, actin and myosin run often perpendicular to the image plane and can be discerned as small and medium sized puncta. This more clearly reveals the organization of the acto-myosin bundling and holds the potential for further analysis of their packing (Figure 4d).

In order to assess the quality of the data acquired, we picked 33,000 80S ribosome particles from 200 randomly selected tomograms from the double-sided attachment experiment. Subtomogram averaging and classification yielded a structure at a resolution of 6.9 Å (GSFSC, Figure 5a). The resolution was likely limited by lamella thickness, supported by the fact that the CTF could not be fit beyond 6 Å.

**Figure 5:**
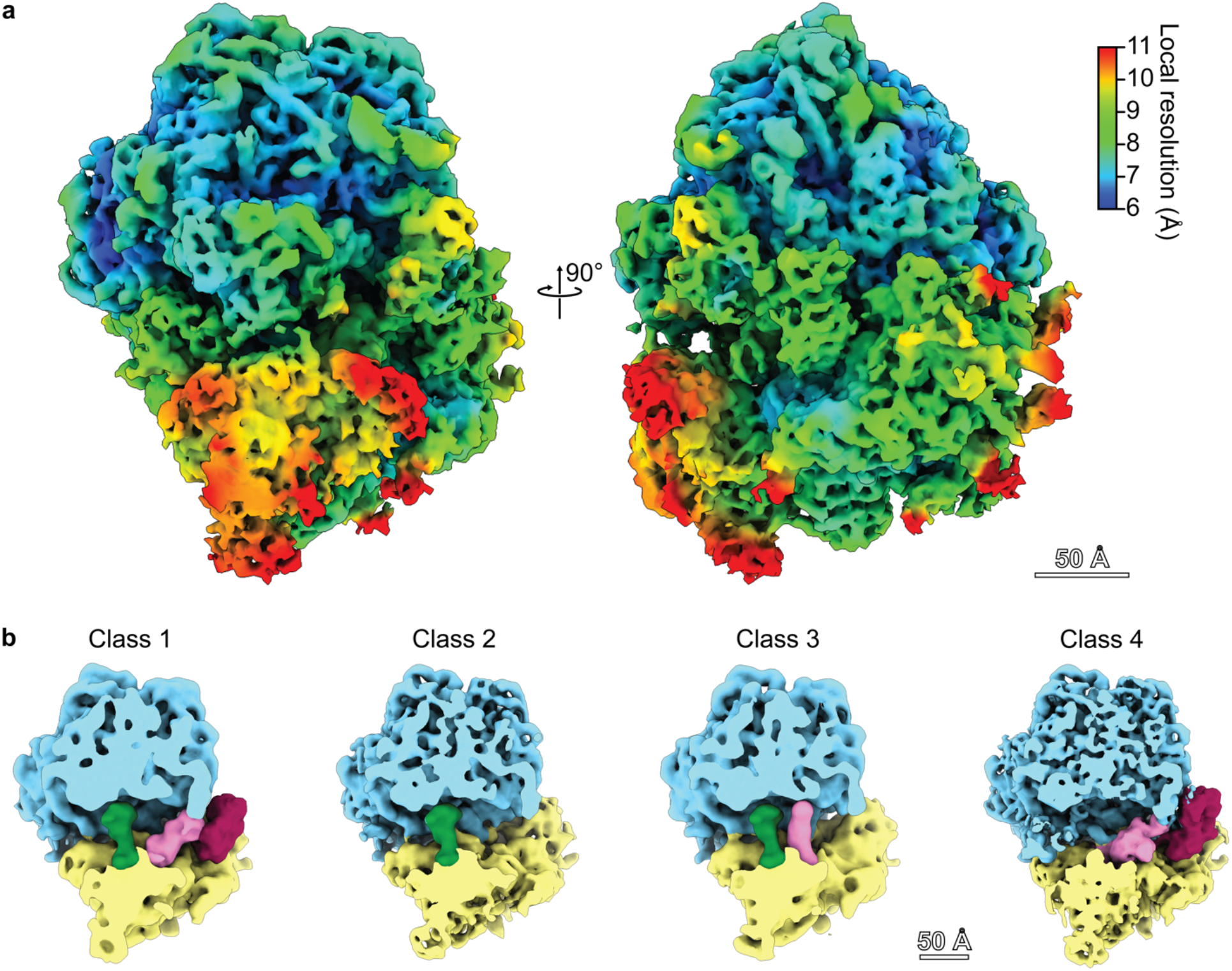
Subtomogram average reconstruction of the *C. elegans* 80S ribosome from *in situ* data. **a**, The *C. elegans* ribosome to a resolution of 6.9 Å (GSFSC). Density map is colored by local resolution. Note that protein α-helices and rRNA helices are clearly visible at this resolution. **b**, Four different ribosomal states obtained through subtomogram classification: Ribosome Class 1 with occupied A, P and EF sites, Ribosome Class 2 with an occupied P site, Ribosome Class 3 with occupied A and P sites, Ribosome Class with occupied A and EF sites.

Classification yielded four different sub-populations in various translational states. When comparing these states to the recently published ribosome state landscape from *D. discoideum* [8], resemblances are apparent to the initiation state with an occupied P site (Figure 5b – Class 2), states with an occupied A and P site (Figure 5b – Class 3) and elongation factor bound states (Figure 5b – Class 1, Figure 5b – Class 4).

## Discussion

In the past, cryo-lift-out has been hampered by limited throughput and low overall success rate. Therefore, the technique has even been deemed ‘practically impossible’, or at least ‘merely difficult’ in the literature (Parmenter et al 2016). The fact that the existing literature has not advanced beyond proof-of-principle experiments underlines this evaluation. The study of multicellular organisms and tissues by cryo-electron tomography, however, holds an enormous potential for biological discovery and technical advances are thus needed. We anticipate the combination of recent hardware and workflow improvements (Klumpe et al 2022, Parmenter & Nizamudeen 2021, Schreiber et al 2018) with the increased throughput of Serial Lift-Out to make cryo-ET data acquisition from high-pressure frozen material more attainable. In addition, plasma focused ion beam technology, while available for some years, was recently introduced to the cryo-FIB community (Berger et al 2023, Martynowycz et al 2023, Sergey et al 2018) and may further improve cryo-lift-out throughput by increased ablation rates.

The cryo-FIB lamella milling protocols developed to date remove most of the cell during lamella preparation. The final lamella represents only <1 % of a eukaryotic cell and even less for multicellular organisms. While a high number of lamellae milled at different heights could restore that lost information through an ensemble average (Ferguson et al 2017), Serial Lift-Out lamellae originate from a single specimen, yielding a more thorough characterization of its morphology. A variety of volume microscopy methods, e.g. volume electron microscopy or X-Ray tomography, retain this information. They remain, however, limited in resolution when compared to cryo-ET. Serial Lift-Out increases the contextual information retained in cryo-ET while preserving data acquisition at molecular resolution.

Here, we have shown the ability to section in increments of one to four micrometers. This process has two steps that contribute to specimen loss: sectioning (∼300-500 nm) and lamella milling. The latter can be minimized by finer sectioning. Preliminary experiments suggest that thinner sectioning down to 500 nm may be achievable. Reducing the section thickness even further in order to prepare lamellae directly from the extracted volume, though, will likely require major technological advances of the FIB instrument. Developments in ion beam shaping could become valuable for such endeavors, reducing the material lost during sectioning. Due to the physical basis of ablation in FIB preparation, however, material will always be lost to the milling process itself. Therefore, truly consecutive lamellae, similar to serial sections of plastic embedded samples, seem unlikely to be attainable.

Nevertheless, the creation of multiple sections within biological material increases the preparation throughput for HPF samples and adds contextual information. For model organisms with a stereotypic and well-described body plan such as *C. elegans* or *Platynereis dumerilii (Vergara et al 2021)*, Serial Lift-Out sections and tomograms may be mapped back into context using other sources of volumetric data as illustrated in Figure 3a (Britz et al 2021, Dumoux et al 2023). As a result, back-mapping analysis may enable label-free targeting of features and events that are tissue and cell-type specific.

Serial Lift-Out also addresses the challenge that arises when dealing with anisotropic cells and tissues. As the sectioning plane of the specimen can be adapted, fluorescence or FIB/SEM data can inform the preparation of the lift-out site and, in turn, the sectioning angle. This adaptive preparation strategy can give new insights into the molecular architecture as illustrated by tomograms from transverse sections of muscle cells. When compared to previously obtained longitudinal sections (Burbaum et al 2021, Davide et al 2023, Wang et al 2021), the transverse section shows actin-myosin packing from a novel angle revealing how actin filaments surround what is likely to be myosin heads. The combination of acquiring cryo-ET data on both the transverse and longitudinal sections also hold the potential to improve subtomogram analysis when facing structures with preferential orientation.

In addition to guiding site preparation for lift-out, cryo-correlative light and electron microscopy is more generally used to target specific subcellular events (Arnold et al 2016, Bieber et al 2022, Klumpe et al 2021, Schaffer et al 2019). This technique, however, remains challenging for routine use within larger tissues. Improvements in the operation of integrated light microscopes are therefore necessary to streamline subcellular targeting in cryo-lift-out experiments. Serial Lift-Out combined with in-chamber light microscopy could increase the success rate of targeting by reducing the sample thickness used in the correlation, increasing the number of sections and in turn targeting attempts, and enabling to regularly check the fluorescence signal throughout the milling process.

One limitation of cryo-lift-out is the high rate of ice contamination during transfer. As can be seen in the TEM overviews (Figure 3 – Figure Supplement 1), large ice crystals tend to obstruct regions of the lamellae and prevent data acquisition. Serial Lift-Out, similar to on-grid preparations, compensates for the loss in imageable area through a higher yield of lamellae in comparison to previous lift-out techniques. Reducing transfer contamination, however, remains highly desirable and controlled environments, e.g. glove boxes (Tacke et al 2021), or other technological advances such as vacuum transfers will be needed to maximize the imageable area of cryo-FIB lamellae.

Finally, the analysis of lamella thickness distributions and lamella survival rate suggest that double-sided attachment in cryo-lift-out is advantageous. This like likely due to the reduction of lamella bending during milling and greater stability during the transfer to the TEM.

With the methodological advances of Serial Lift-Out, the existing hurdles of lift-out have been greatly diminished, enabling data quality, throughput, and success rate in cryo-lift-out that permits the mapping of large tissue regions and whole organisms at the molecular level. Tomography on lamellae of an L1 larva obtained with our Serial Lift-Out method elucidated its ribosome to 6.9 Å resolution and uncovered four different translational states. Taken together, Serial Lift-Out demonstrates enormous potential to discern and study the molecular anatomy of native tissues and small multicellular organisms.

## Supporting information

Figure2_MovieSupplement1

Figure2_MovieSupplement2

Figure3_MovieSupplement1

## Acknowledgement

We thank Jonathan Wagner, Adriana Prajica, Philipp S. Erdmann, Johann Brenner, Meijing Li, David Klebl, Alexander Rigort, Bernhard Hampoelz, Martin Beck, and Reinhard Fässler for invaluable input and support. We thank the Caenorhabditis Genetics Center and Bloomington Drosophila Stock Center for providing *C. elegans* and *D. melanogaster* strains, respectively. This study used the infrastructure of the Department of Cell and Virus Structure at the MPI of Biochemistry. SK was supported by the International Max Planck Research School for Molecular and Cellular Life Sciences. JMP acknowledges funding from the Max-Planck Society.

## Data availability

Tomograms have been deposited in the Electron Microscopy Data Bank (EMDB) under accession codes: EMD-17246, EMD-17247, EMD-17248, Subtomogram averages have been uploaded under accession numbers: EMD-17241, EMD-17242, EMD-17243, EMD-17244, EMD-17245 and will be released upon publication.

## Materials and Methods

### Sample Vitrification

*C. elegans* strains AM140 (allele rmIs132[unc-54p::Q35::YFP]) and NK2476 (allele qy46[ina-1::mKate+loxP]) were cultivated according to standard methods on rich NGM (Brenner 1974). To obtain a synchronous population of animals, gravid young adult worms were washed from five petri dishes (6 cm) with M9 medium. The worms were pooled into 15 mL centrifuge tubes and centrifuged at 175x g. Excess supernatant was removed and the worm suspension was mixed with bleach solution (3.75 mL 1M NaOH, 3.0 mL household bleach, 8.25 mL H_2_O (Sulston & Hodgkin 1988) and swiftly vortexed. Bleaching was continued with intermittent vortexing and checking on a stereomicroscope. As soon as roughly 50% of worms appeared broken, the bleaching procedure was stopped by the addition of egg-buffer (118 mM NaCl, 48 mM KCl, 2 mM CaCl2, 2 mM MgCl2, 25 mM HEPES, pH 7.3) and centrifugation at 175g for 1 minute. The supernatant was swiftly removed and the egg/worm carcass solution was washed another four times with egg-buffer. The egg/worm carcass mixture was then floated on 60 % sucrose solution and purified eggs were withdrawn from the surface (Strange et al 2007). The egg solution was washed in M9 buffer and larvae were allowed to hatch at 20 °C for 24 hours.

This population of developmentally arrested L1 larvae was used for vitrification, carried out following a modified version of Mahamid, *et al*. (Mahamid et al 2015) on a Leica EM-ICE (Leica Microsystems, Wetzlar, Germany). Sample carriers Type B (Leica Microsystems, Wetzlar, Germany) with a diameter of 6 mm were coated with a separating layer of cetyl palmitate solution (0.5% w/v in diethylether) by dipping the carriers briefly into the solution and placing them, cavity side down, on filter papers to let the solvent evaporate. A 2 µL drop of 20% (v/v) Ficoll 400 im M9 was applied to the flat side of the sample carrier and a Formvar-coated grid (50 mesh square or 75 mesh hexagonal mesh) was floated on this drop with the support film facing the liquid. Any excess cryoprotectant below the grid was wicked away, using pieces of Whatman No 1 filter paper (Whatman, Maidstone, UK). Next, synchronized L1 larvae were mixed with an equal amount of 40% Ficoll 400 in M9 medium to reach a final concentration of 20% cryoprotectant. 2µL of this sample solution was applied onto the grid and the sandwich was completed by addition of a second 6 mm sample carrier Type B, cavity-side up. The sample was immediately high-pressure-frozen. Grids were removed from the HPF sample carriers and stored in LN_2_ until further use. EM-grids were clipped into Thermo Fisher Scientific cartridges for FIB-milling. To be able to locate the biological material (L1 larvae) in the ice layer, clipped grids were either previously mapped in a Leica SP8 confocal microscope equipped with a cryostage or directly transferred into a FIB-SEM instrument with an integrated light microscope.

*Drosophila* samples were prepared as previously described (Klumpe et al 2021). In brief, fly strains were maintained at 22 °C on standard cornmeal agar. 24 hours prior to dissection of egg chambers, the flies were transferred into a new vial supplemented with yeast paste. Egg chambers were dissected in Schneider’s medium. For high-pressure freezing, 3 mm HPF carriers were soaked in hexadecene and blotted on filter paper. Depending on the egg chamber stage, the dissected material was subsequently transferred into the 100 µm or 150 µm cavity of a 3 mm Type A or Type C HPF sample carrier (Engineering office M. Wohlwend, Sennwald, Switzerland), respectively. The filling medium was 20% Ficoll (70 kDa) in Schneider’s medium. The flat surface of a Type B HPF Planchette was used to close the HPF assembly for freezing in the Leica EM ICE high-pressure freezer (Leica Microsystems, Wetzlar, Germany). The sample was subsequently pre-trimmed using a 45° diamond knife (Diatome, Nidau, Switzerland) in a cryo-microtome (EM UC6/FC6 cryo-microtome, Leica Microsystems) at a temperature of -170 °C and used for further FIB preparation.

### Cryo-fluorescence microscopy

Grids were clipped in Thermo Fisher Scientific (TFS) cartridges and imaged using a Leica TCS SP8 laser confocal microscope (Leica Microsystems, Wetzlar, Germany) fitted with a Leica cryo-stage (Schorb et al 2017). Imaging was performed with a 50x 0.90 NA objective and HyD detectors.

Tile set montages of the NK2476 strain sample were collected using a pinhole size of 4.85 AU and a voxel size of 0.578 µm × 0.578 µm × 2.998 µm. The tile sets were merged and maximum intensity projections for each channel calculated using LAS X (Leica, Wetzlar, Germany). Autofluorescence imaging was performed for the double-sided attachment experiment. The 488 nm laser line at 2.5 % total power was used for excitation and the emission was detected from 501 nm to 535 nm. The reflection channel was excited at 552 nm and 2% total laser power and detected with an emission range of 547 nm to 557 nm.

Tile set montages of the AM140 strain were collected at a pinhole size of 1 AU and a voxel size of 0.578 µm × 0.578 µm × 1.03 µm. The tile sets were merged and maximum intensity projections for each channel were calculated using LAS X (Leica, Wetzlar, Germany). Q35-YFP was excited at 488 nm and 0.5 % total laser power and emission was detected from 509 nm to 551 nm. Reflection was recorded with 638 nm excitation at 0.83 % total laser power and the emission range set to 633 nm to 643 nm.

### Serial Lift Out - Single Side Attachment

#### Lift-In Grid Preparation

100 square mesh Cu grids (Agar Scientific, Stansted, UK) were assembled into TFS cartridges. The cartridges were marked for manual alignment during sample loading. These markings were aligned to where the grid bars intersect the cartridge marks. The marks were also used for grid orientation during loading. The grids were then loaded into a FIB-SEM 45° pre-tilt cryo-shuttle such that the grid bars are aligned vertically and horizontally.

Once loaded, 14-pin grids were prepared by rotating the grid plane to be normal to the ion beam (trench milling orientation) and milling out two combs of horizontal grid bars. This yielded grids with 14 pins that can be used for 28 lamella attachments (Figure 2 - Figure Supplement 2). A horizontal grid bar was used to prepare 20 µm × 10 µm × 5 µm copper blocks for attachment to the lift-out needle by redeposition milling using single-pass regular cross-sections directed away from the needle (Figure 1 – Figure Supplement 2, Figure 2 – Figure Supplement 5a). Note, that the copper block attached to the needle can be re-used several times. The lower section of the copper block used for the previous attachment can be milled away leaving a clean surface for the next attachment.

As the FIB-SEM does not have to be cooled during lift-in grid milling, we recommend the preparation of several lift-in grids ahead of time in order to reduce workload during the serial lift-out session.

#### Lift-Out, Sectioning and Milling Procedure

Serial lift-out was performed on an Aquilos 2 FIB-SEM instrument (Thermo Fisher Scientific, Waltham, MA, U.S.A.) equipped with an EasyLift system. To facilitate lift-out, the EasyLift needle was modified by attaching a copper block adapter (see ‘lift-in grid preparation’) using redeposition. The copper adapter has a higher sputter yield and, thus, increased redeposition compared to tungsten and vitreous ice. Additionally, the block increases the surface area available for attachment to the biology, and reduces the wear of the needle over time.

After copper block attachment, the volume to be extracted was prepared by milling trenches around the region of interest in trench milling orientation. The fluorescence data acquired of the L1 ‘waffle’ samples was correlated to the ion beam images using MAPS version 3.14 (Thermo Fisher Scientific, Waltham, MA, U.S.A.). The grid was coated with a protective metal-organic platinum layer using a gas injection system (GIS) heated to 27°C at a stage working distance of 9.72 mm and a stage tilt of 45° for 90s. Trenches were milled with the grid perpendicular to the ion beam (trench milling orientation) after which the EasyLift needle was inserted. In order to increase the redeposition yield for attachment, the copper block needs to be aligned below the sample surface and flush to the surface exposed by trench milling. Redeposition from the copper block onto the extraction volume was achieved using single-pass patterns of regular cross-sections instead of the default multi-pass mode to avoid re-milling previously redeposited material. These patterns were placed at the interface of the two surfaces on the copper adapter with the milling direction for these patterns directed away from the extraction volume (Figure 2 – Figure Supplement 5b). The extraction volume was then released through milling the last connection to the bulk sample with a line pattern. The 30 µm × 110 µm × 25 µm extracted volume was subsequently lifted from the bulk sample.

After the extraction of the volume, the shuttle was returned to the lamella milling orientation (18° in a system with a 45° pre-tilt) and the stage translated to the receiver grid. The lower edge of the extracted volume and the corner of the pin were aligned in both electron and ion beam images. Attachment was performed by redeposition, using single-pass regular cross-section patterns directed away from the extracted volume (Figure 2 – Figure Supplement 5d). Once attached, the EasyLift system was maneuvered by a 50 nm step horizontally and vertically away from the pin in horizontal direction in order to create a small amount of strain. Subsequently, the lower 4 µm of the volume were sectioned from the rest of the volume using a line pattern. This process was iterated until the entire extracted volume had been sectioned.

### Serial Lift-Out: Double-Sided Attachment

#### Lift-In Grid Preparation

For double-sided attachment, 100/400 rectangular mesh copper grids (Gilder, Grantham, UK) were used as the receiver grid. These were clipped into standard TFS cartridges that marked such that the 400 mesh bars were in line with the bottom mark and the 100 mesh bars were in line with the side markings. This grid was loaded into the FIB shuttle and 20 µm × 10 µm × 5 µm copper block adapters were prepared as described above.

During the preparation of the receiver grid, roughly 5 µm deep line patterns were milled across the horizontal 400 mesh grid bars in trench milling orientation to divide the rectangles approximately ⅓ and ⅔. These marker lines are intended to guide the alignment of the extracted volume by providing a reference visible in both the electron and ion beams during the attachment process.

#### Lift Out, Sectioning and Milling Procedure

Lift-out site preparation for double-sided attachment was performed on an Aquilos 1 FIB-SEM instrument (Thermo Fisher Scientific, Waltham, MA, U.S.A.) equipped with a METEOR in-chamber fluorescence light microscope (Delmic, Delft, Netherlands)(Smeets et al 2021). The the GIS was employed for 90 s at 27°C with at a stage working distance set to 10.6 mm and the grid normal to the GIS needle to apply a layer of protective metal-organic platinum. The METEOR in-chamber microscope was used in conjunction with previously acquired cryo-confocal data to localize the larva of interest and produce an extraction volume of 42 µm × 180 µm × 20 µm. A width of 40 µm is necessary in order to span the space between the grid bars of a 100/400 rectangular mesh grid. A slight excess of width of the extraction volume is preferable, since a too narrow block will not allow for double-sided attachment, while material can be ablated during sectioning.

The lift-out and receiver grid were subsequently transferred into an Aquilos 2 FIB-SEM instrument (Thermo Fisher Scientific, Waltham, MA, U.S.A.) equipped with an EasyLift system. Lift-out was otherwise performed as stated in the section for single-sided attachment. In brief, the volume was attached to the needle via the copper adapter, the volume was released, the needle retracted, and the shuttle was returned to the lamella milling orientation at the receiver grid. The extracted volume was re-inserted, and its lower edge was positioned in-between two grid bars. In the case when the volume was slightly too wide, material was ablated from its sides as necessary. To allow for proper attachment, the volume’s lower edge was aligned to the reference line on the receiver grid in both, the electron and ion beam images and the volume was attached on both sides by redeposition from the grid bars. To achieve this, single-pass cross-section patterns were placed on the grid bars above the marker lines in in close proximity of the interface between the grid bars and extracted volume, while the milling was directed away from the volume (Figure 2 – Figure Supplement 5c). Once attached, the EasyLift system was maneuvered up in the z-direction by 50-100 nm in order to create a small amount of strain. Subsequently, a section was released from the extracted volume by milling a line pattern at 4 µm above the volume’s lower edge. This process was iterated until the entire extracted volume had been sectioned.

**Table 1:** Milling pattern parameters for Serial Lift-Out. The table summarizes the parameters used for the milling steps necessary to perform a Serial Lift-Out experiment. The corresponding pattern geometries are illustrated in Figure 2 – Figure Supplement 5. All patterns were milled at 30.00 kV FIB acceleration voltage. Pattern files for Thermo Fisher Scientific instruments are provided in Figure 2 – Pattern Files Supplement. These templates and the milling parameters may need adjustment for different projects and for FIB-SEM instruments from other manufacturers, specifically concerning differing scanning strategies deployed by the microscope manufacturer.

| Step | Pattern Type | Beam Current | Multipass Number | Approximate Time |
| --- | --- | --- | --- | --- |
| Copper block milling | Regular Cross-section | 5-65 nA | 4 | 20 mins |
| Copper block attachment | Regular Cross-section | 1 nA | 1 | 3 mins |
| Trench milling | Regular Cross-section | 3 nA | 4 | N/A |
| Extraction volume attachment | Regular Cross-section | 1 nA | 1 | 3 mins |
| Extraction volume attachment | Line | 3 nA | N/A | 10 mins |
| Pin or grid bar attachment | Regular Cross-section | 1 nA | 1 | 4-8 mins |
| Sectioning | Line | 0.5-1 nA | N/A | 8-12 mins |

#### Fine milling of lift-out sections

For both single-sided and double-sided attachment, the initial 4 µm sections were thinned in two steps: rough milling and fine milling. Rough milling was performed using regular cross-section patterns at a beam current of 1 nA to a thickness of 1.5 µm, 0.5 nA to 1.2 µm lamella thickness, 0.3 nA to 0.8 µm lamella thickness. After rough milling all of the sections, fine milling was performed using regular rectangle patterns at a beam current of 0.1 nA to 0.4 µm and 50 pA to final thickness. Over-tilting and under-tilting by up to 1° was used for beam convergence compensation. The double-sided attached lamellae were sputter coated with platinum after fine milling for 4 seconds, at a chamber pressure of 0.20 mbar and a current of 15.0 mA.

#### Serial lift-out from high-pressure frozen *D. melanogaster* egg chambers

Lift-out experiments of high-pressure frozen *D. melanogaster* egg chambers were performed on a Scios FIB-SEM instrument (Thermo Fisher Scientific, Waltham, MA, U.S.A.) equipped with an EasyLift needle. The preparation of the lift-out volume as described previously (Klumpe et al 2021). In brief, an approximately 20 µm × 20 µm volume of target material was prepared using regular cross-section patterns in trench milling orientation in a horseshoe-like shape. The trenches were ∼20 µm wide, except for the region that needed to be accessed by the lift-out needle, which was ∼35 µm wide. The extraction volume was undercut at a stage tilt of 45°, or the highest tilt reachable for the specific position. This preparation leaves the extraction volume attached to the bulk material on a single side. Subsequently, the procedure for single-sided attachment was performed as described above. Sectioning was performed in increments of 1-3 µm using a line pattern milled at a beam current of at 0.3 nA. After section preparation, fine milling was performed by sequentially decreasing the beam current: 0.3 nA to 800 nm, 0.1 nA to 500 nm, 50 pA to 350 nm and 30 pA to < 300 nm. The final step was judged by the loss of contrast in SEM imaging at 3 kV acceleration voltage and a beam current of 13 pA.

#### Lamella preparation by the ‘waffle’ milling method

For preparation of lamellae directly on the high-pressure frozen grid, vitrified as described above, we followed a similar workflow as published as the ‘waffle’ method (Kelley et al 2022). The grid was coated with a metal-organic platinum layer by GIS deposition for 20 s at a stage working distance of 10.6 mm. Milling was performed on a gallium FIB-SEM Aquilos 2 (Thermo Fisher Scientific, Waltham, MA, U.S.A.) instrument. Initial trenches were milled in trench milling orientation at a beam current of 3 nA using regular cross-section patterns. To avoid damaging the region of interest by milling at high beam currents, 2 µm of buffer distance were left around the region of interest. Trenches were extended to 15 µm on the backside of the subsequent lamella and 30 µm on the front side. In a next step, the trenches were extended to 1 µm closer to the region of interest at a beam current of 1 nA and in a last step the final dimension of the lamella region was defined milling at 0.5 nA. After trench milling, another layer of protective metal-organic platinum was deposited on the sample (20 s deposition time, 10.6 mm stage working distance, three consecutive times). The stage was rotated into lamella milling orientation and a notch was milled with line patterns at 0.3 nA as previously described (Kelley et al 2022). The preparation of the final lamella was carried out by removing material above and below of the region of interest at a beam current of 0.3 nA, until a final lamella thickness of 2 µm was reached. In sequential steps of decreasing beam current (to 0.8 µm thickness at 0.1 nA beam current, to 0.4 µm at 50 pA), the remnant material was ablated. The lamella was polished to a final thickness of about 200-250 nm at 30 pA beam current.

#### TEM Data Acquisition

TEM data acquisition was performed on a Titan Krios G4 at 300 kV equipped with a Selectris X energy filter and a Falcon 4i camera (Thermo Fisher Scientific, Eindhoven, The Netherlands). Lamella overview montages were recorded by stage-driven tiling at 11,500x nominal magnification (pixel size 2.127 nm). Tomograms were recorded using the Tomo5 software package version 5.12.0 (Thermo Fisher Scientific, Eindhoven, The Netherlands) using the EER file format.

Two acquisition strategies were deployed. For the tomograms of the single-sided attachment and ‘waffle’ preparation tilt series were acquired at a nominal magnification of 42,000x resulting in a pixel size at the sample of 2.93 Å. A dose-symmetric tilt scheme was used with an angular increment of 2°, a dose of 2 e^-^/Å^2^ per tilt and a target defocus of -4 to -5.5 µm. Tilt series were collected in a tilt range of -70° to 50° due to the lamellae pre-tilt and with a total dose of 120 e^-^/Å^2^.

For the tomograms collected of the double-sided attachment lamellae, that were subsequently used for subtomogram analysis, tilt series were acquired at a nominal magnification of 64,000x, resulting in a pixel size of 1.89 Å. Data was collected using a dose-symmetric tilt scheme with an angular increment of 3°, a dose of 3.23 e^-^/Å^2^ per tilt and a target defocus range of -1 to -4 µm. Angles of -70° to 50° were acquired, resulting in a total dose of 132 e^-^/Å^2^.

#### Tomogram Reconstruction, Visualization and Subtomogram Analysis

Data was processed using the Tomoman version 0.7 pipeline (https://github.com/williamnwan/TOMOMAN). 14 frames with a dose of 0.23 e^-^/Å^2^ per frame were rendered from the EER files. These were used for motion correction in MotionCor2 version 1.4.7 (Zheng et al 2017) and CTF estimation with CTFFIND4 version 4.14 (Rohou & Grigorieff 2015). Bad tilts were removed after manual inspection using the Tomoman script. Dose-weighting was performed at 3.23 e^-^/Å^2^ per tilt using either Tomoman or Warp. For denoising, tilt series were separated into odd and even tilts during motion correction and the resulting stacks were processed using Cryo-CARE (Buchholz et al 2019). Tomogram reconstructions for visualization were done in IMOD version 4.12.32. Tilt series were aligned with AreTomo version 1.3.3. CTF-corrected tomograms for template matching were reconstructed in IMOD version 4.12.32 (Kremer et al 1996, Mastronarde & Held 2017) at eight times binning, resulting in a pixel size of 15.6 Å.

Initial template matching was performed in STOPGAP version 0.7 (Wan et al 2020) on a subset of 70 tomograms at bin8 using PDB 4V4B as a reference filtered to 35 Å (Spahn et al 2004). 16,420 particles were extracted and aligned in STOPGAP to generate the *C. elegans* 80S ribosome template. The template was subsequently used to repeat template matching on 200 tomograms at bin8. 65,451 particles were extracted and cleaned by projecting the subtomogram along the Z-axis and subsequent 2D classification in Relion version 4.0. The remaining data set contained 37,026 particles. The retained particles were reprocessed in Warp version 1.0.9, cleaned to remove particles with inadequate CTF resolution and astigmatism. The resulting 35,350 particles were extracted with a pixel size of 2.98 Å and a boxsize of 160 pixels. Relion version 3.0 with a spherical mask of 340 Å radius was used to align the subtomograms. Finally, the particles were imported into M version 1.0.9 and geometric and CTF parameters were sequentially refined. Corrected subtomograms were extracted from M and classified in Relion 3.0. Two rounds of classification with a spherical mask of 340 Å in diameter resulted in 8,256 particles being removed. The second step was a focused classification with a spherical mask around the A/P/E-site. The resulting 5 classes were combined into 2 classes according to the small subunit rotation. Each of these 2 merged classes were subjected to another round of 3D classification with a mask around the elongation factor binding site. The final classes were manually pooled according to structural similarity yielding 5 classes with 2,175, 20,638, 877, 1,119, and 2,285 particles, of which the class with 877 particles was neglected. Segmentation of the elongation factor and tRNAs was done with the Segger tool in Chimera. ChimeraX version 1.3 was used for visualization (Pettersen et al 2021).

