## Extended_Data_File for "Serial Lift-Out – Sampling the Molecular Anatomy of Whole Organisms"

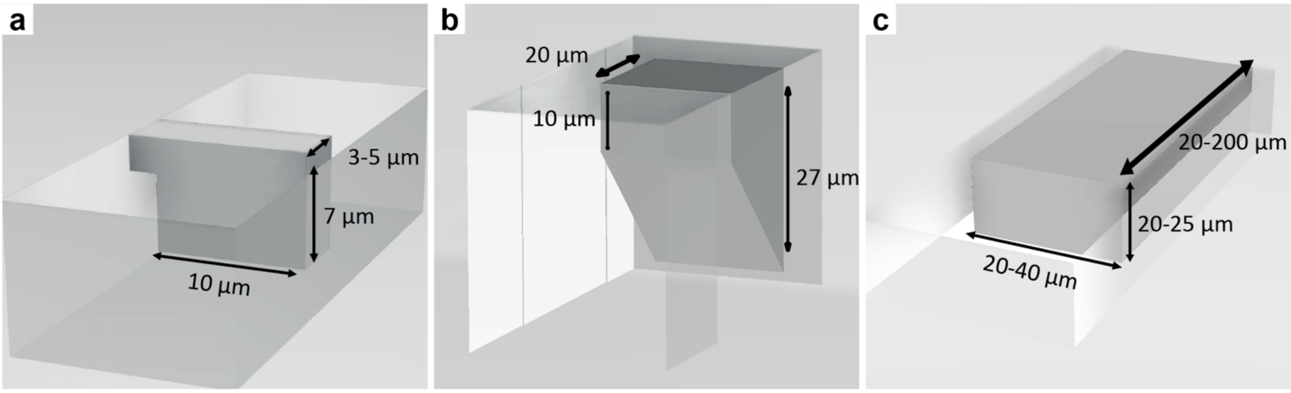


**Figure 1 – Figure Supplement 1: Schematics of lift-out geometries.** Geometries of the extracted volume for different lift-out approaches are illustrated. Extraction volumes are depicted in dark grey, with the empty volume around them indicating trenches milled prior to the lift-out procedure. **a**, Lamella lift-out prepares a thin section with a small amount of excess material around the final lamella volume. **b**, Lift-out from voluminous HPF samples requires detachment of the material from the bulk by an undercut. This lift-out geometry for thick HPF preparations generally limits the extraction volume to approximately 20-30 µm in height, dependent on the angle of the undercut. **c**, Lift-out from ‘waffle’-type samples allows the creation of large extraction volumes. As no undercut is required, the entire thickness of the sample can be extracted.

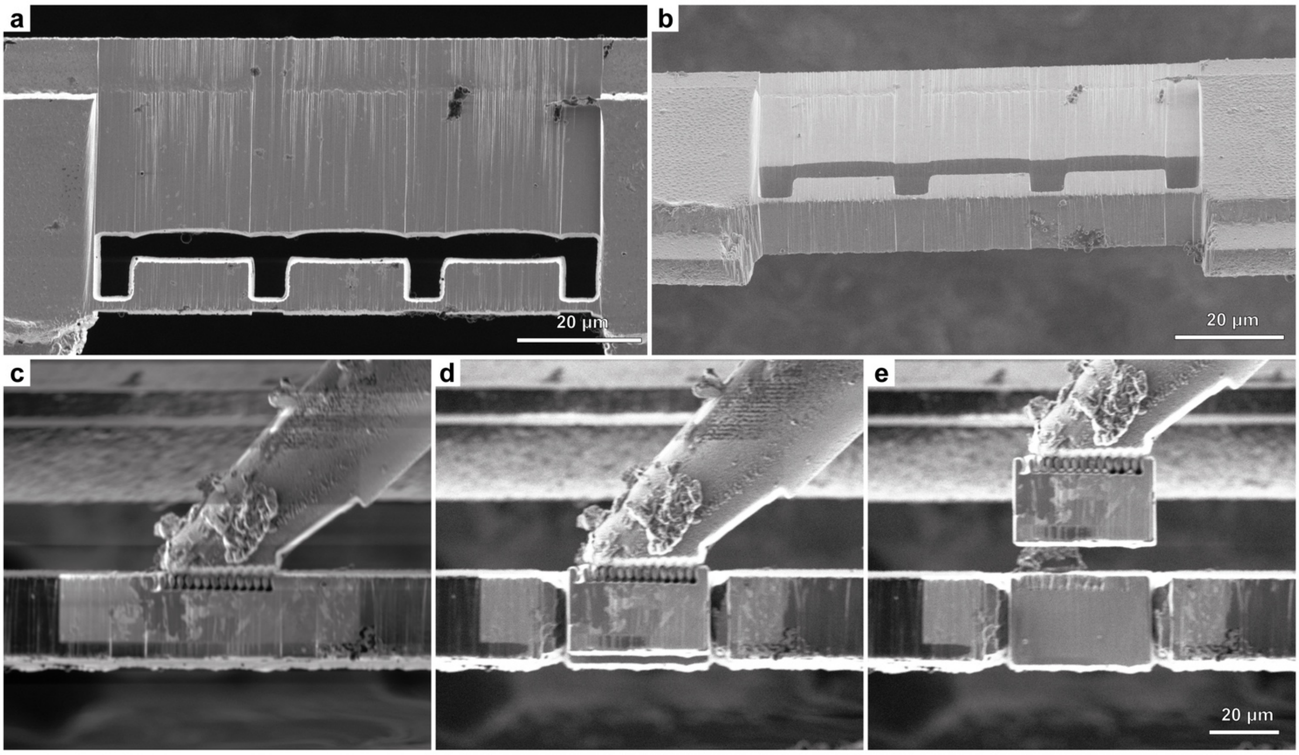


**Figure 1 – Figure Supplement 2: Copper block adapter preparation and attachment.** The grid bars of the receiver grid can be used to prepare copper blocks that serve as adapters for the attachment of the extracted volume to the tungsten needle. **a**, FIB top view and **b**, SEM side view of a grid bar, with copper blocks prepared for extraction. Three blocks were milled into a bar of a copper 100 mesh grid (trench milling orientation). **c-e**, FIB images at lamella milling orientation. The tungsten needle of the micromanipulator was flattened to increase the copper block contact area and the block is attached by redeposition milling (c). The copper block is subsequently milled free from the grid bar (**d**) and lifted-out (**e**). All steps were performed using a 45° pre-tilt shuttle.

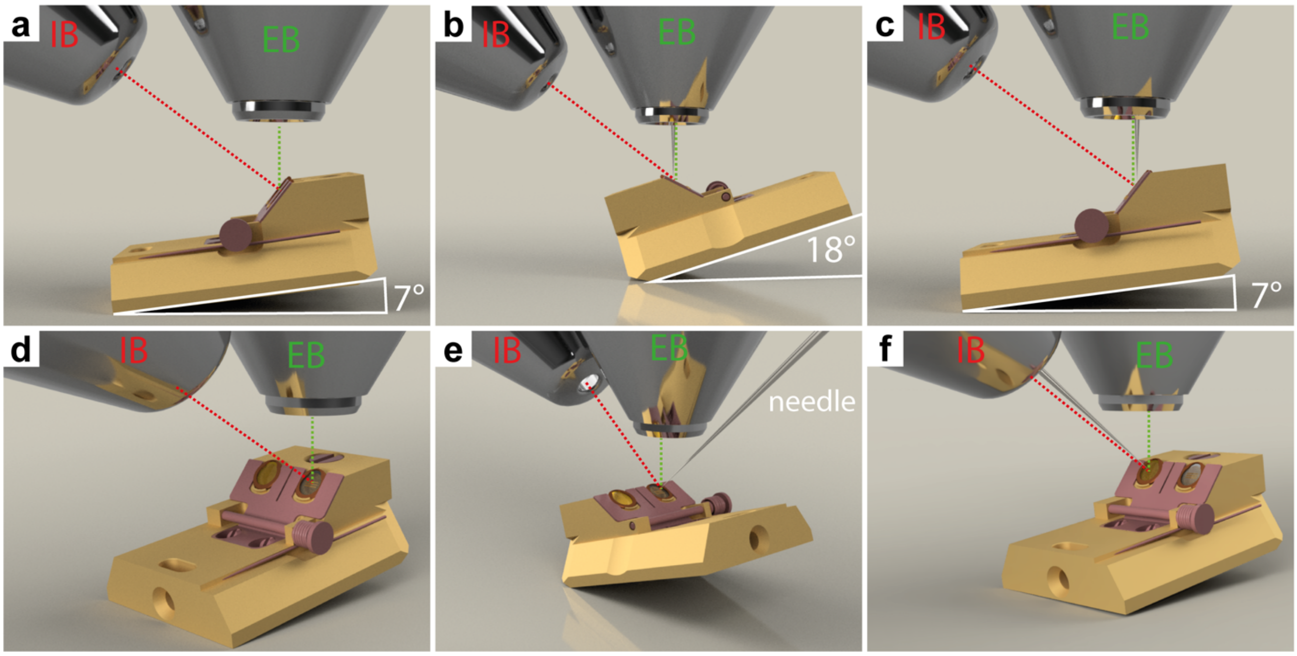


**Figure 1 - Figure Supplement 3: Stage orientations for Serial Lift-Out. a**,**d**, Copper block preparation and trench milling are performed at 7° tilt angle and 180° relative stage rotation ('trench milling orientation'). **b**,**e**, Lift-out of the copper block is performed at 18° tilt angle and 0° relative rotation ('lamella milling orientation'). The same orientation is used during the attachment of Serial Lift-Out slices to the receiver grid, lamella thinning the common lift-out procedure from voluminous HPF samples. **c**,**f**, Lift-out to obtain sections perpendicular to the grid plane is performed at trench milling orientation. Panels **d**-**f** are oblique front views of the respective side views shown in **a**-**c**. From this perspective, the HPF sample is located in the left shuttle position and the receiver grid is located the right shuttle position. Panels **b**,**c**,**e**,**f** show the EasyLift-needle inserted. EB and IB indicate the column of the ion beam and electron beam, respectively.

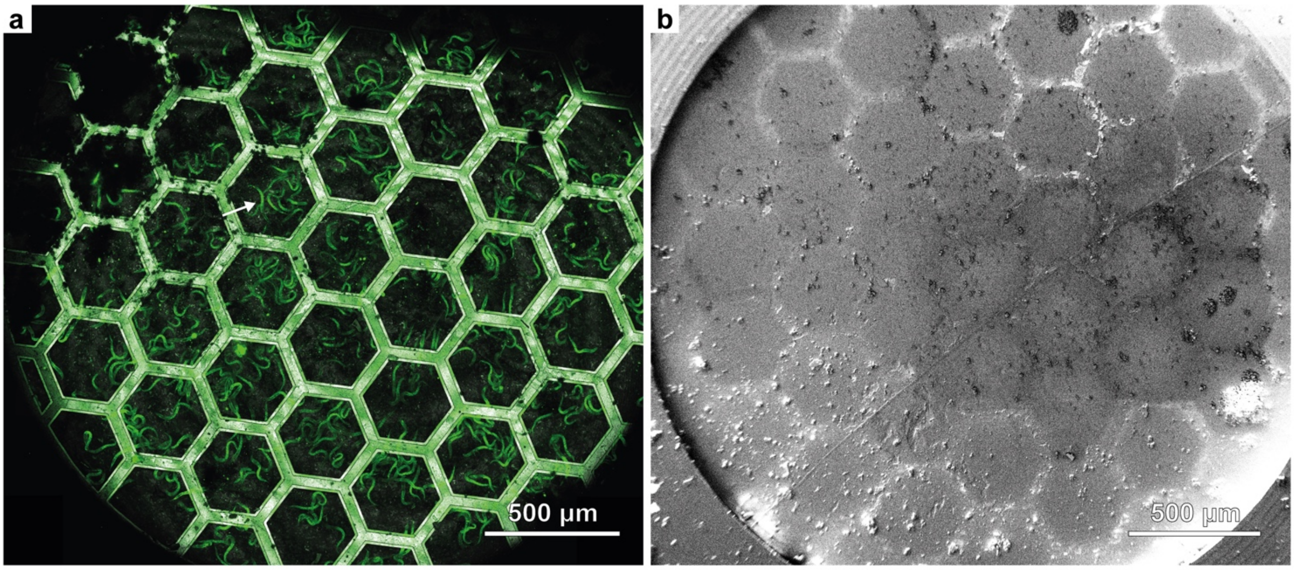


**Figure 2 - Figure Supplement 1: Overview of the ‘waffle’-type grid used for the double-sided attachment Serial Lift-Out. a**, Maximum-intensity projection of the fluorescence (green) and reflected light (grey) channels **b**, SEM overview of the grid shown in (**a**).

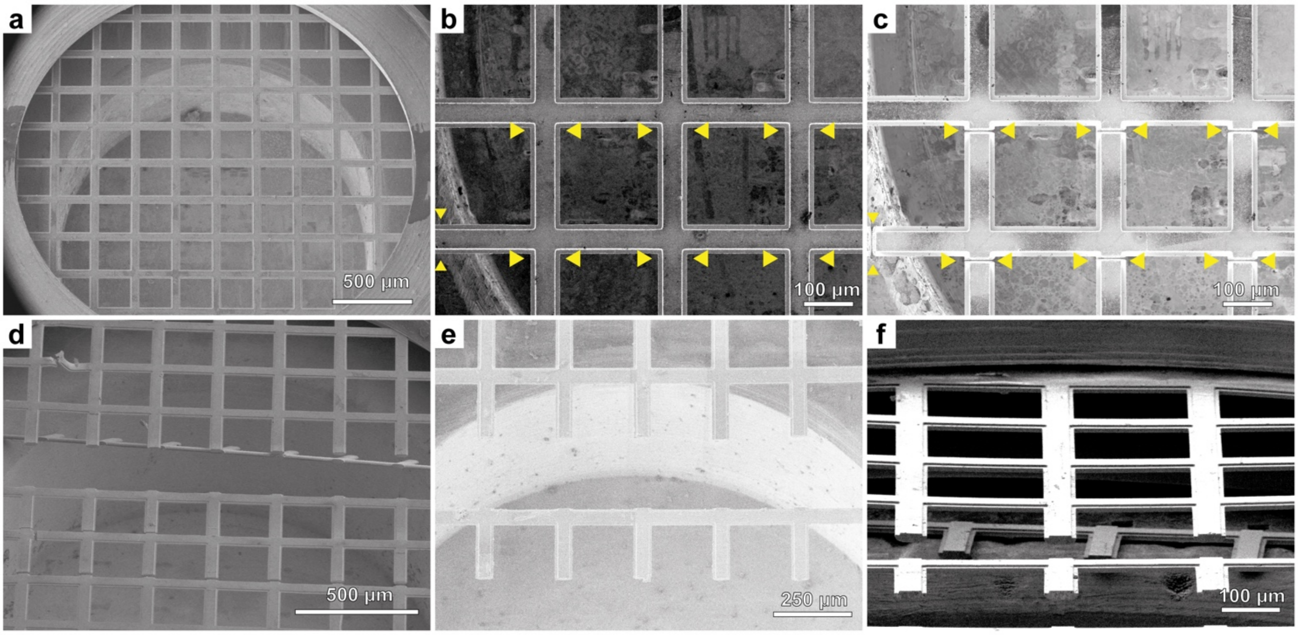


**Figure 2 - Figure Supplement 2: Preparation of a Serial Lift-Out receiver grid for single-sided attachment.** **a**, SEM image view of a 100 square mesh copper grid clipped into an AutoGrid. **b**,**c**, FIB images of grid bars prior to (**b**) and after (**c**) milling line patterns (between yellow arrowheads, trench milling orientation, beam current 65 nA). **d**, SEM image of the first grid bar after milling completion. **e**,**f**, SEM (**e**) and FIB (**f**) view of the final grid after both grid bars were removed (lamella milling orientation).

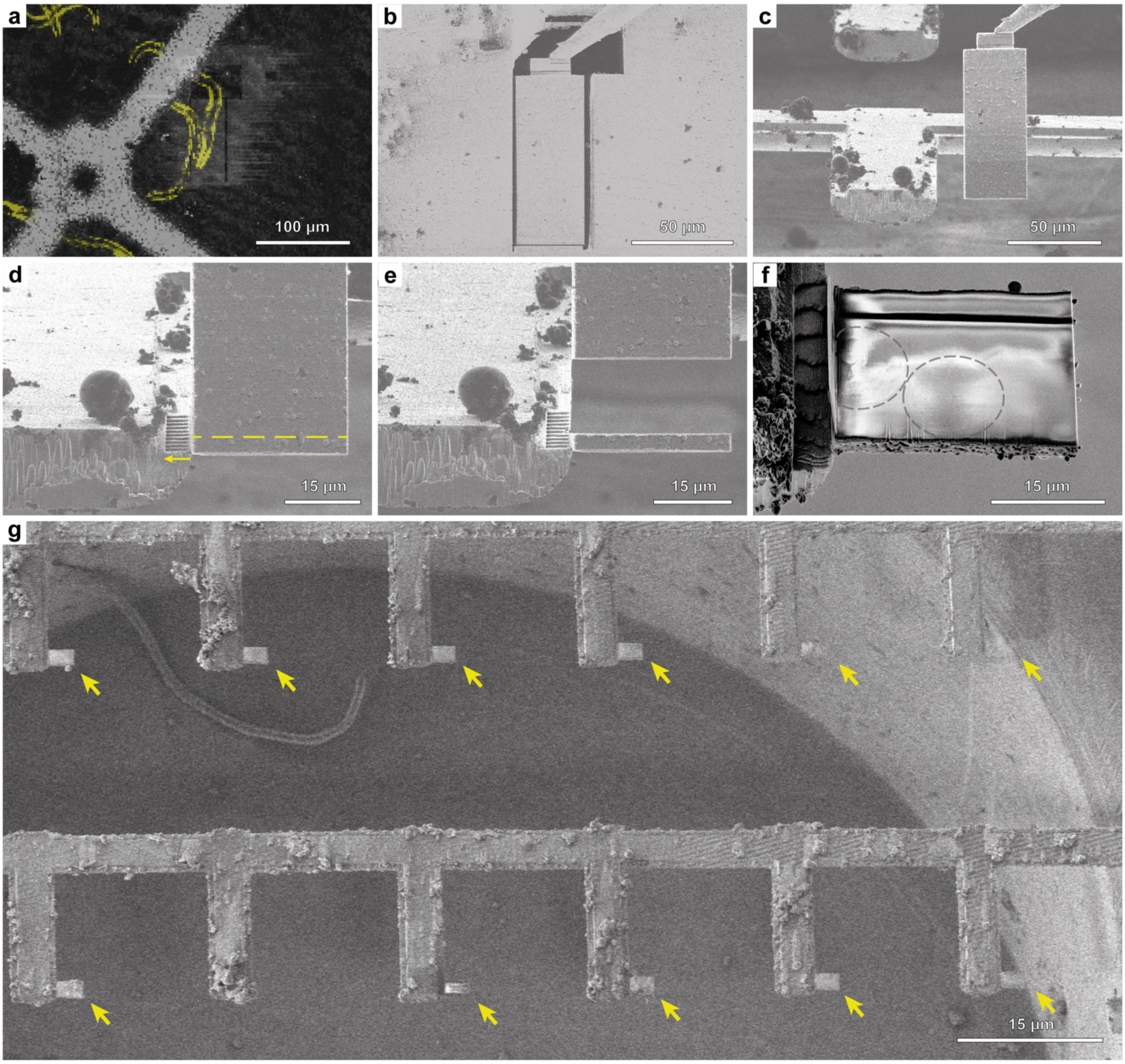


**Figure 2 – Figure Supplement 3: A workflow for single-sided attachment Serial Lift-Out. a**, FIB image of the extraction site with overlaid correlated fluorescence data (yellow) indicating the larva being targeted (trench milling orientation). **b**, The extraction volume is attached to the EasyLift needle using redeposition from the copper adapter. The release cut, milled with a line patten, is noticeable at the base of the extraction volume (trench milling orientation). **c**, FIB image of the extracted volume being lowered to the attachment position adjacent to a pin. For attachment, the lower front edge of the volume is aligned to the corner of the pin. **d**, Attachment using redeposition from the pin (yellow arrow indicates milling direction), followed by line pattern milling releasing the section of a desired thickness (dashed yellow line). **e**, The resulting section is depicted as the remaining extracted volume is retracted. **f**, SEM image of a typical section after release from the extraction volume. Note the faint pattern of worm cross-sections discernible (grey dashed lines). **g**, SEM image of the resulting receiver grid after a session of single-sided attachment Serial Lift-Out. Arrows indicate the 12 sections obtained. Figure 2 - Movie Supplement 2 summarizes the process.

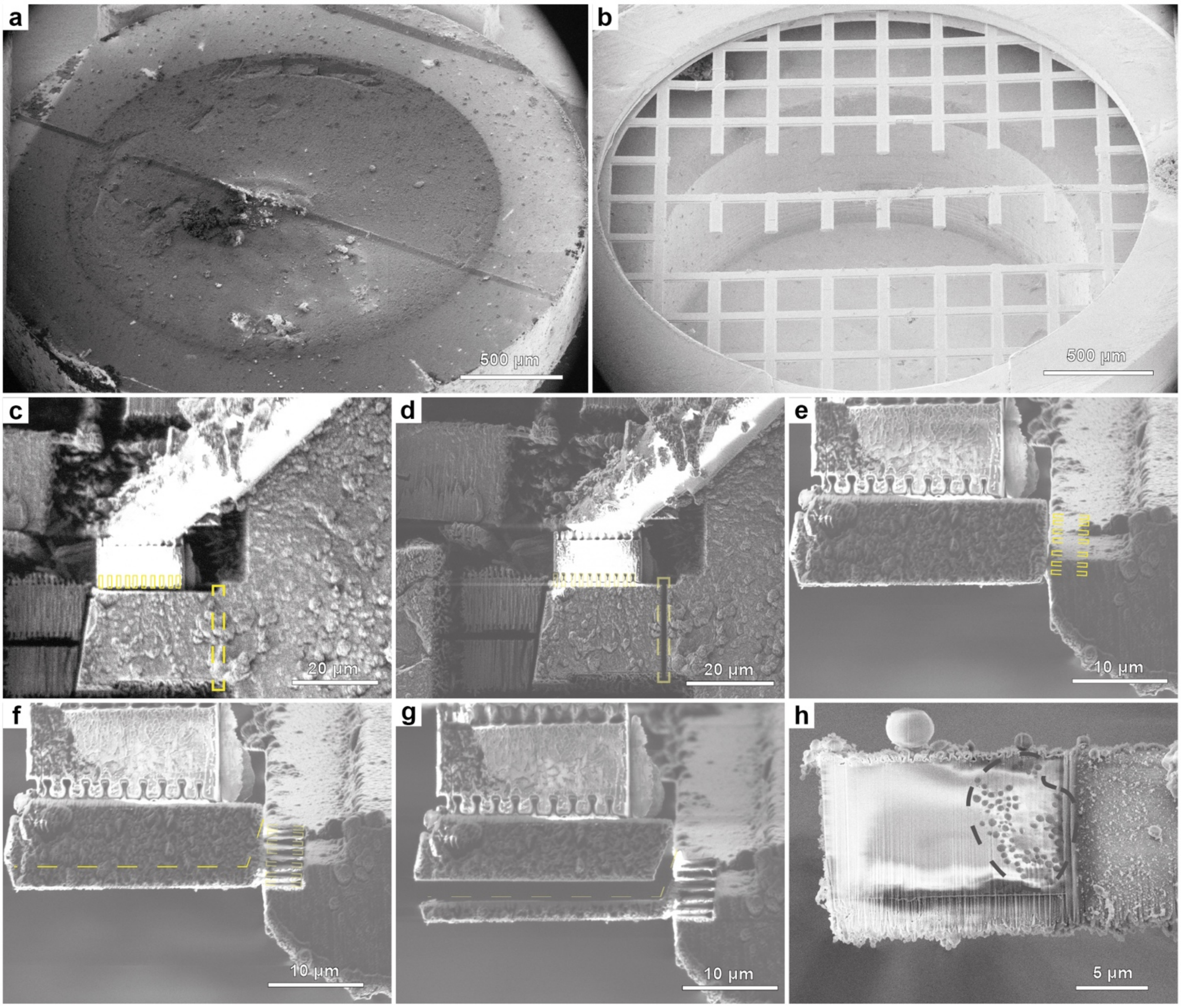


**Figure 2 - Figure Supplement 4: Voluminous HPF samples by single-sided attachment Serial Lift-Out.** Serial Lift-Out on *D. melanogaster* egg chambers vitrified in HPF sample carriers. **a**, HPF sample carrier planed by diamond knife trimming at 45° in a cryo- ultramicrotome to access regions deep within the egg chamber. **b**, Receiver grid as prepared by FIB milling (Figure 2 – Figure Supplement 2). **c**, FIB image of the target extraction volume before attachment needle system. Redeposition patterns are indicated with yellow boxes. **d**, The extracted volume is released from the bulk by milling a regular cross-section pattern (dashed yellow box) and transferred to the receiver grid. **e**, FIB image of the extracted volume being lowered to the attachment position adjacent to a pin. For attachment, the lower front edge of the volume is aligned to the corner of the pin. Redeposition patterns are indicated with yellow boxes. **f**, The extracted volume is attached to the pin by redeposition from the grid bars (yellow dashed boxes in **e**). The section is separated from the remaining volume by line pattern milling (dashed line). **g**, The remaining volume is retracted for successive rounds of sectioning. The sliced section remains attached to the pin. **h**, SEM image view of the section shown in **g** after polishing the surface by low current milling. Note the clearly visible lipid droplets in the cytoplasmic region of the egg chamber nurse cell (grey dashed line). The region to the left, lacking lipid droplets, is the nucleoplasmic region that was targeted prior by fluorescence microscopy as confirmed by subsequent TEM imaging. **c-d,** Trench milling (perpendicular to ion beam) orientation. **e-h**, Lamella milling orientation.

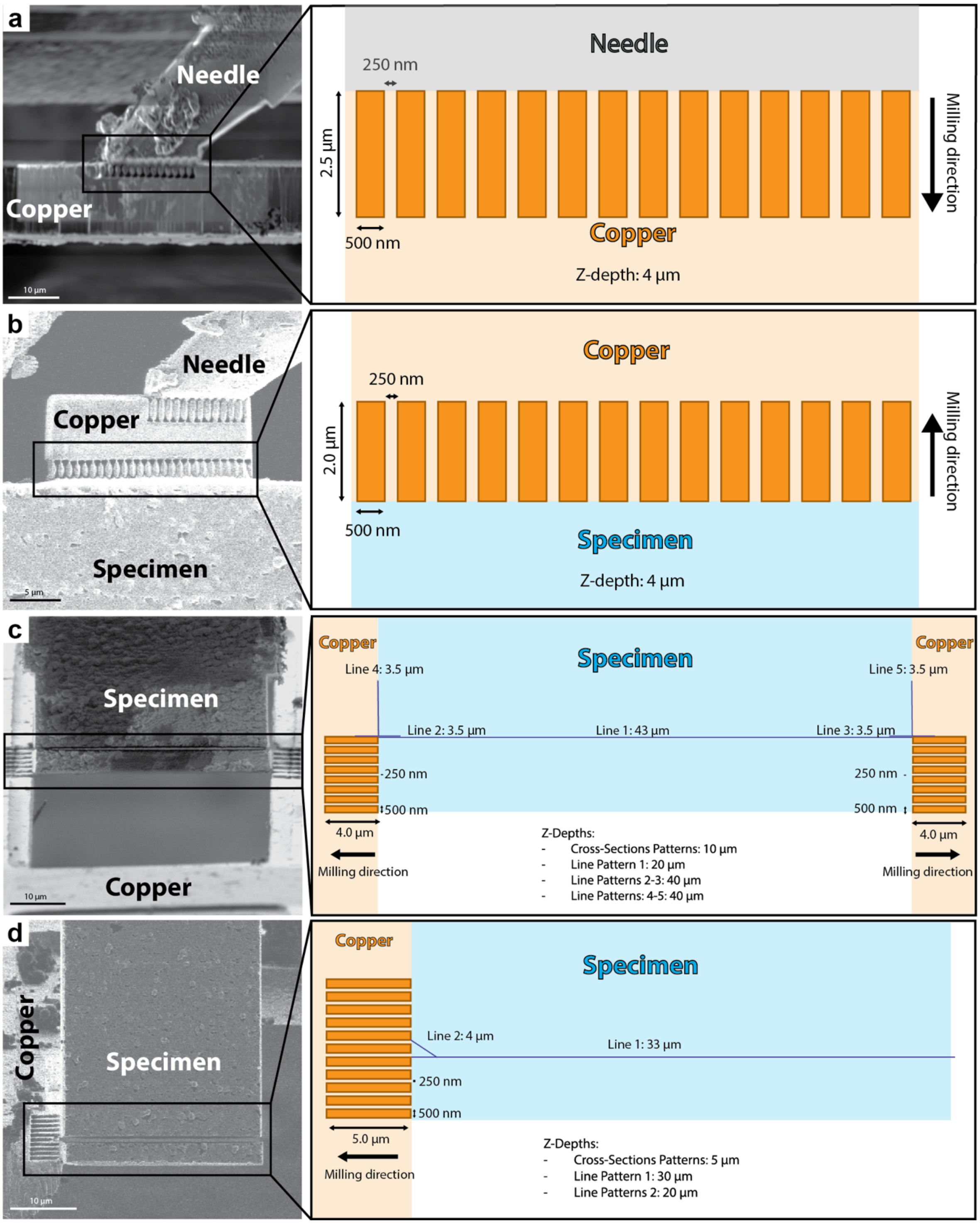


**Figure 2 – Figure Supplement 5: Serial Lift-Out milling pattern dimensions.** A schematic of the milling patterns used for double-sided and single-sided attachment Serial Lift-Out. Orange rectangles represent single-pass cross-section patterns used for attachment, while blue lines indicate line patterns used to sectioning. Note that the exact dimensions of the redeposition patterns may need to be adjusted for different instruments. **a**, Copper block redeposition attachment to the EasyLift system. **b**, Copper block redeposition attachment to the extracted volume. **c**, Double-sided attachment: Redeposition attachment from the grid bars to the specimen, followed by section release by line pattern milling. **d**, Single-sided attachment: Redeposition attachment from the pin to the specimen, followed by section release by line pattern milling. Pattern files for Thermo Fisher Scientific FIB-SEM instruments are available for download in Figure 2 – Pattern Files Supplement.

**Figure 2 – Movie Supplement 1: Double-sided Serial Lift-Out workflow.** A movie summarizing the Serial Lift-Out process with double-sided attachment. The L1 larva was targeted by correlating a fluorescence overview to the grid surface. Lift-out is performed from the trench milling orientation. Copper redeposition is used to attach the adapter to the extraction volume. The volume is released by milling a line pattern and the EasyLift needle is retracted. The stage is set to lamella milling orientation on the receiver grid, and the needle is reinserted. The extracted volume's lower edge is aligned to a previously milled line mark (line pattern across the bars) and redeposition from the grid bars is used to attach the volume. The bottom section is released from the remaining extracted volume using line pattern milling. This process was repeated in order to create 40 serial lift-out sections.

**Figure 2 – Movie Supplement 2: Single-sided Serial Lift-Out workflow.** A movie summarizing the Serial Lift-Out process with single-sided attachment. The L1 larva was targeted by correlating a fluorescence overview to the grid surface. Lift-out is performed from the trench milling orientation. Copper redeposition is used to attach the adapter to the extraction volume. The volume is released by milling a line pattern and the EasyLift needle is retracted. The stage is set to lamella milling orientation on the receiver grid, and the needle is reinserted. The extracted volume's lower edge is aligned to a corner of the pin and redeposition from the pin is used to attach the volume. The bottom section is released from the remaining extracted volume using line pattern milling. This process was repeated in order to create 12 serial lift-out sections.

**Figure 2 - Pattern Files Supplement.** ThermoFisher Scientific FIB-SEM instrument pattern files for various Serial Lift-Out steps: 1. Attachment of the copper block to the needle (Block_to_needle_glue.ptf), 2. Attachment of the copper block to the extraction volume (Block_to_bio_glue.ptf), 3. Double-sided attachment using a 100/400 rectangular mesh copper grid (Double_sided_attachment.ptf) and 4. Single-sided attachment using pins generated from a 100 square mesh copper grid (Single_sided_attachment.ptf). Schematics of the patterns are shown in Figure 2 – Figure Supplement 5.

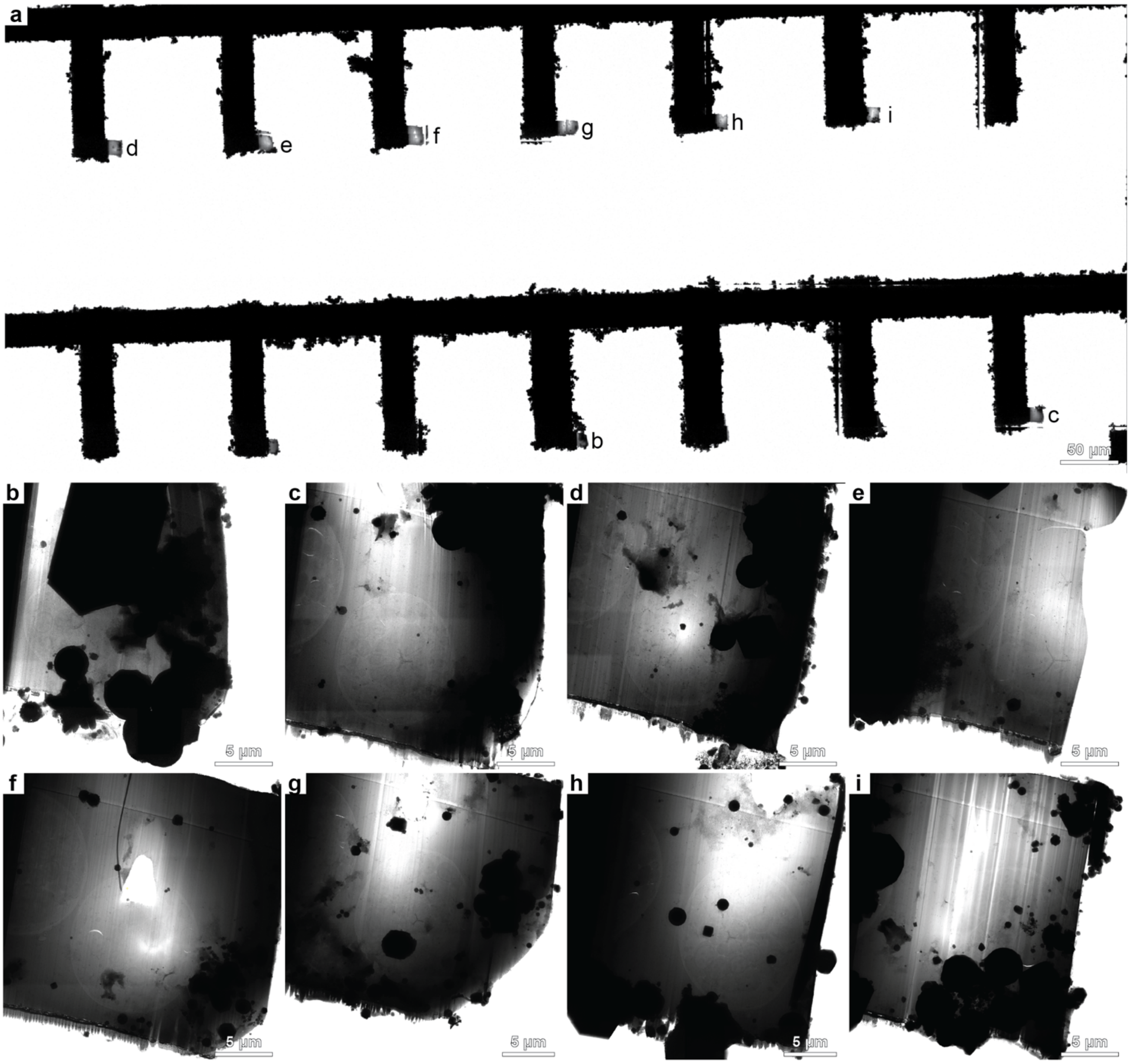


**Figure 3 – Figure Supplement 1: TEM overview of the grid derived from single-sided attachment Serial Lift-Out.** **a**, TEM low magnification overview image (magnification 125x) of the lamella region of the Serial Lift-Out receiver grid. **b-h**, Lamella montage overviews (magnification 11,500x) of 8 out of 12 fine-milled lamellae that survived transfer into the TEM and on which tilt series were acquired. The letters in panel **a** indicate the corresponding higher magnification micrographs in panels **b**-**h**.

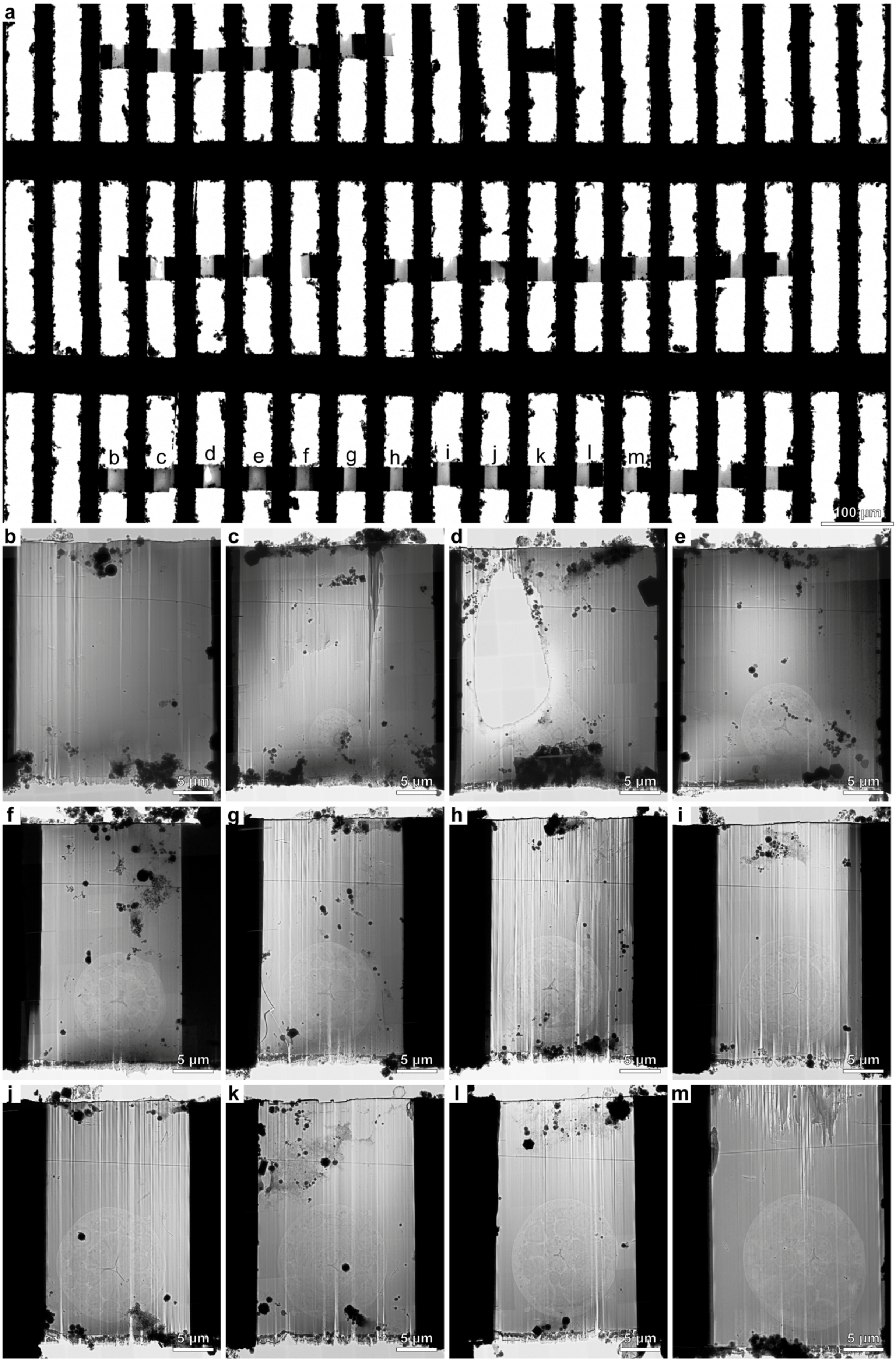


170

**Figure 3 – Figure Supplement 2: TEM overview of the grid derived from the double-sided attachment Serial Lift-Out experiment. a**, TEM low magnification overview image (magnification 125x) of the lamella region of the Serial Lift-Out receiver grid for double-sided attachment. **b**-**m**, Lamella overviews (magnification 11,500x) of the first 12 sections. In total, 40 sections were prepared, of which 39 were milled to lamella thickness. 33 were successfully transferred into the TEM. The letters in panel **a** indicate the corresponding higher magnification micrographs in panels **b**-**m**.

**Figure 3 – Movie Supplement 1: Morphological detail on intermediate magnification overview maps.** The movie illustrates the amount of detail, that can be extracted from lamella overview montages recorded at 11,500x magnification. Cross-sections obtained from the double-sided attachment Serial-Lift-Out experiment are shown. The first section stems from the procorpus, the anterior pharynx region (Figure 3 – Figure Supplement 2e). In the subsequent sections (Figure 3 – Figure Supplement 2f-j), the central pharynx widens up to form the anterior pharyngeal bulb or metacorpus. The seventh section in the movie was taken at the anterior pharyngeal isthmus and contains the anterior dorsal part of the nerve ring. Here, a zoomed in view of the nerve ring illustrates the discernible detail. Section 8 of the movie (Figure 3 – Figure Supplement 2l) is successive to the previous. The camera zooms in on the pharynx, the posterior ventral part of the nerve ring, a golgi apparatus, body wall muscle cells, a lateral amphid process bundle and the seam cell.

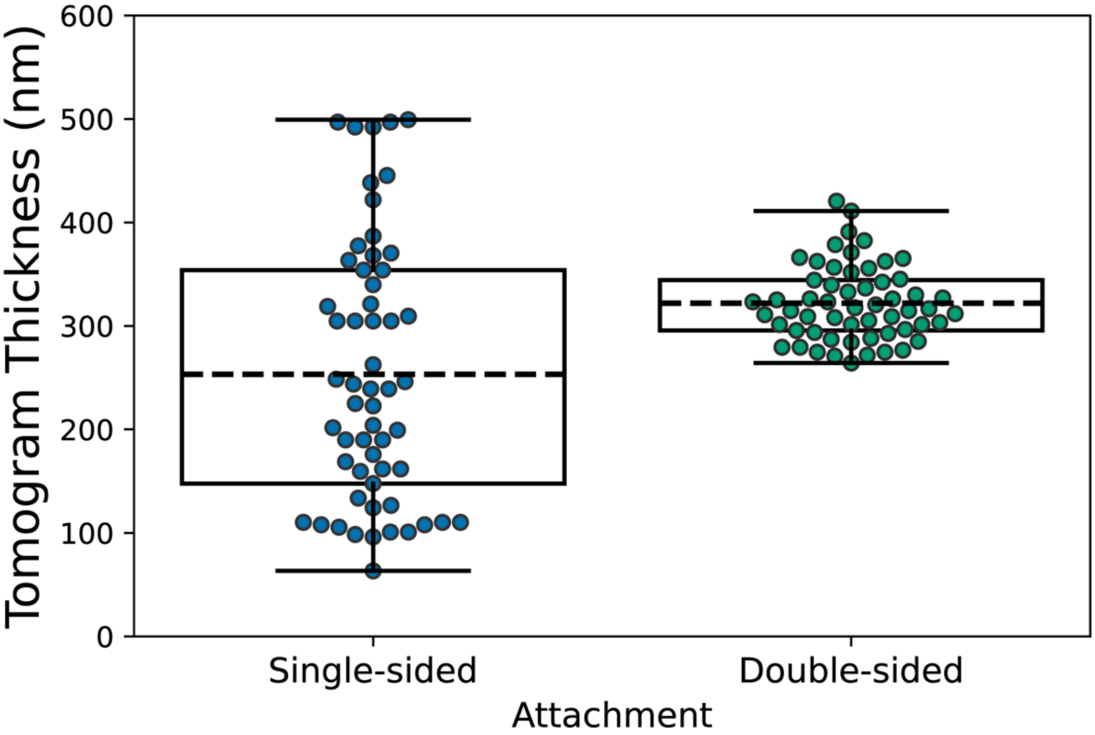


**Figure 4 – Figure Supplement 1: Tomogram thickness distribution for the lift-out experiments.** A box plot for the distribution of thickness measurements on single-sided and double-sided attachment Serial Lift-Out lamellae all 57 tomograms recorded for the single-sided attachment experiment (blue data points) and a random selection of 132 tomograms from the 1012 tilt series collected for the double-sided attachment experiment (green data points). While the average tomogram thickness is lower for single-sided attachment (253 nm ± 125 nm) than for double-sided attachment (303 nm ± 40 nm), the spread of thickness measurements is higher for single-sided attachment, likely due to increased bending of the free-standing lamellae during milling. The box indicates the two middle quartiles of the data, the dashed line the mean of all measurements, the whiskers indicate the interquartile range of the data.

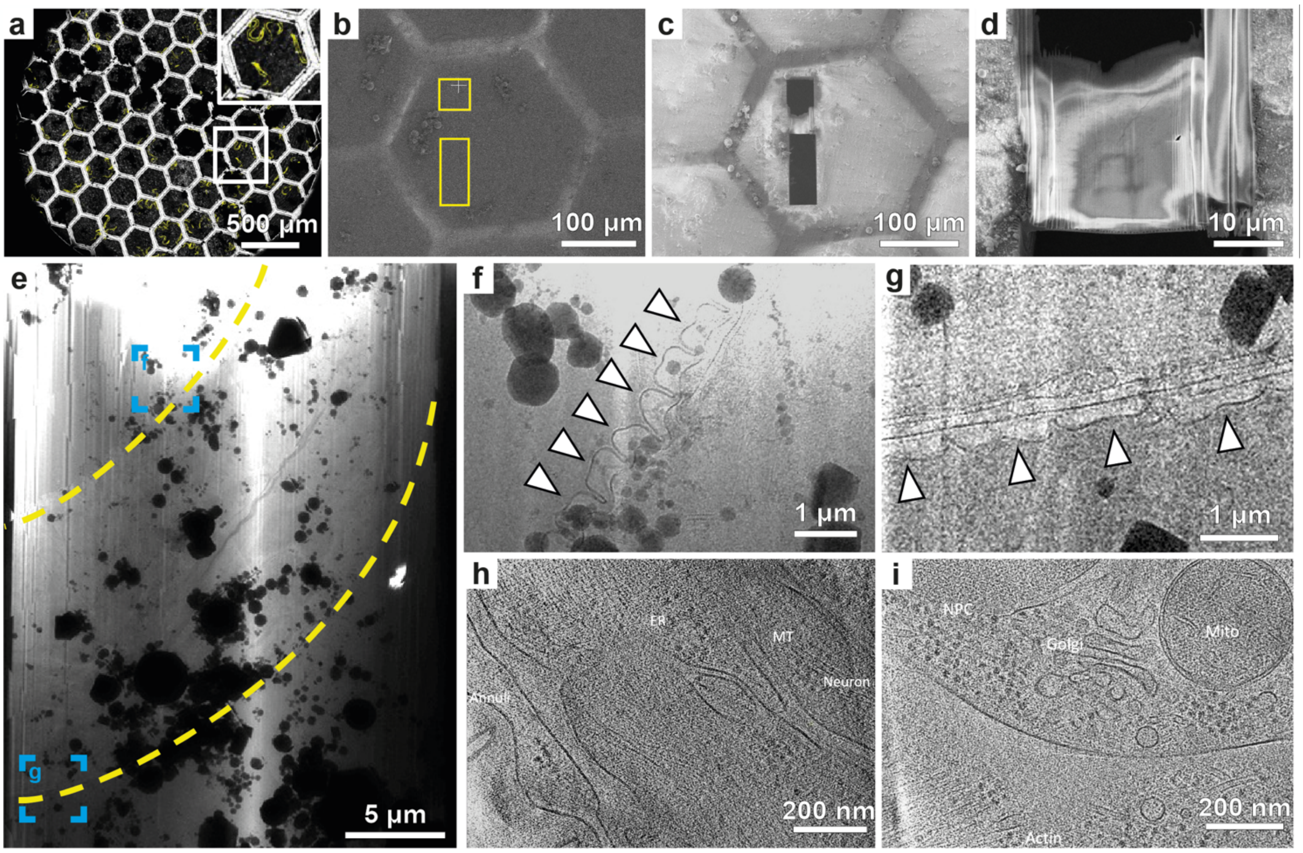


**Figure 4 – Figure Supplement 2: 'Waffle' lamella milling of a *C. elegans* L1 larva. a**, Fluorescence (yellow) and reflection (white) channel confocal micrograph of the grid used for 'waffle' milling. The inset highlights the mesh in which the larva was targeted. **b**,**c**, FIB view images of the mesh shown in inset **a** pre- (**b**) and post-milling (**c**). Trenches were milled at the trench milling orientation (sample surface perpendicular to the ion beam). Yellow rectangles indicate the trench milling locations. **d**, SEM overview of the final lamella. **e**, TEM overview of the lamella. Yellow dashed lines indicate the lateral outline of the larva. **f**,**g**, Structural difference between the body wall regions of the contracted and relaxed sides of the worm freely thrashing before vitrification. This difference can be clearly seen through the morphological change in the annuli (arrowheads). **h**,**i**, Tomograms recorded on the lamella shown in **e** revealing biological features as obtained from the longitudinal section. ER: endoplasmic reticulum, MT: microtubule, NPC: nuclear pore complex, Mito: mitochondria.

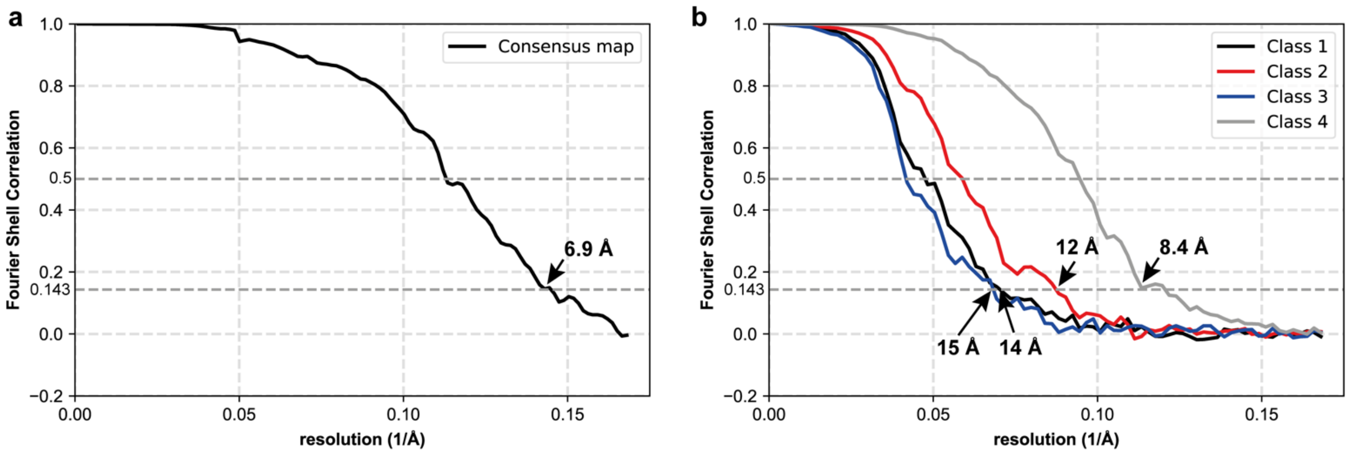


**Figure 5 – Figure Supplement 1**: **a** FSC curves of the consensus average and **b** the four different translational states of the *C. elegans* ribosome
